# Multiple Brain Activation Patterns for the Same Perceptual Decision-Making Task

**DOI:** 10.1101/2023.04.08.536107

**Authors:** Johan Nakuci, Jiwon Yeon, Nadia Haddara, Ji-Hyun Kim, Sung-Phil Kim, Dobromir Rahnev

**Affiliations:** School of Psychology, Georgia Institute of Technology, Atlanta, Georgia, 30332, USA; Department of Psychology, Stanford University, Stanford, California, 94305, USA; Department of Biomedical Engineering, Ulsan National Institute of Science and Technology, Ulsan, South Korea

**Keywords:** perceptual decision making, clustering, fMRI, default mode network

## Abstract

Meaningful variation in internal states that impacts cognition and behavior remains challenging to discover and characterize. Here we leveraged trial-to-trial fluctuations in the brain-wide signal recorded using functional MRI to test if distinct sets of brain regions are activated on different trials when accomplishing the same task. Across three different perceptual decision-making experiments, we estimated the brain activations for each trial. We then clustered the trials based on their similarity using modularity-maximization, a data-driven classification method. In each experiment, we found multiple distinct but stable subtypes of trials, suggesting that the same task can be accomplished in the presence of widely varying brain activation patterns. Surprisingly, in all experiments, one of the subtypes exhibited strong activation in the default mode network, which is typically thought to decrease in activity during tasks that require externally focused attention. The remaining subtypes were characterized by activations in different task-positive areas. The default mode network subtype was characterized by behavioral signatures that were similar to the other subtypes exhibiting activation with task-positive regions. These findings demonstrate that the same perceptual decision-making task is accomplished through multiple brain activation patterns.

## Introduction

Brain activity elicited by the same stimulus or task is highly variable^1,2^. Variation in task-induced brain activity has been identified in individual neurons^3^ and large-scale brain networks^4^ impacting cognition and behavior^5,6^. Variation in brain activity affects our actions in social situations^7^, economic decisions^8^, and even low-level perception^9^.

Despite the widespread variability in brain activity during a task, standard analyses aim to identify the task-induced changes in brain activity across all trials^10^. Such analyses have been applied to a multitude of tasks such as face perception^11^, memory^12^, and navigation^13,14^. The prevailing assumption in studies seeking to identify the brain response to a stimulus or a task is that there is a single pattern of activation. In the case of fMRI, this pattern is typically identified by performing a standard general linear modeling analysis. Under this assumption, trial-to-trial variation in brain activity is simply noise. When applied to tasks that require *externally focused* attention, this standard analysis has identified a set of regions – termed “task-positive” – that increase in activity, and another set of regions – termed “task-negative” – that decrease in activity in response to external demands^15^.

However, it is also possible that subsets of trials can produce meaningfully different patterns of activations that are not well captured by averaging across all trials. Indeed, the blood-oxygen level-dependent (BOLD) signal in fMRI is both spatially and temporally variable^16^, with at least some of this variability likely to stem from meaningful variations in internal processing rather than simply noise^5,6,17,18^. Further, it has been hypothesized that a cognitive process can be accomplished through multiple pathways because of degeneracy^19^, and supported by work utilizing theoretical models and patient populations^20^, but these multiple pathways have not been explicitly identified in healthy individuals.

Here, we sought to identify discrete patterns of brain activity associated with the completion of the same task. To do so, we utilized a data-driven classification method to identify unique patterns in brain activity among individual trials. Across three perceptual-decision making experiments, we found multiple different activation patterns, with one of them surprisingly exhibiting task-negative activations. We further established the behavioral profile associated with each subtype. Finally, we replicated the existence of multiple activations patterns in a working memory task. Overall, our results indicate that multiple brain activation patterns can co-exist in the context of the same task.

## Results

### Variation in brain activity across individual trials

We examined the patterns of brain activation produced across three perceptual decision-making tasks (Experiments 1-3; **Table 1**). We first performed standard GLM analyses to identify task-related brain activations. In Experiment 1, we observed increased activity in the visual and motor cortices, as well as decreased brain activity in the orbital frontal cortex and in the posterior cingulate cortex (**Fig. 1A**). However, single-trial beta responses estimated with a general linear model (GLM) using GLMsingle^21^ deviated substantially from the group map. For example, unlike the group map, trial 2 for subject 1 showed strong positive activity in both the posterior cingulate cortex and the orbitofrontal cortex (**Fig. 1B**). On the other hand, trial 6 for subject 1 produced an activation pattern similar to the group map with negative activity in both the posterior cingulate cortex and the orbitofrontal cortex (**Fig. 1C**).

**Figure 1.**
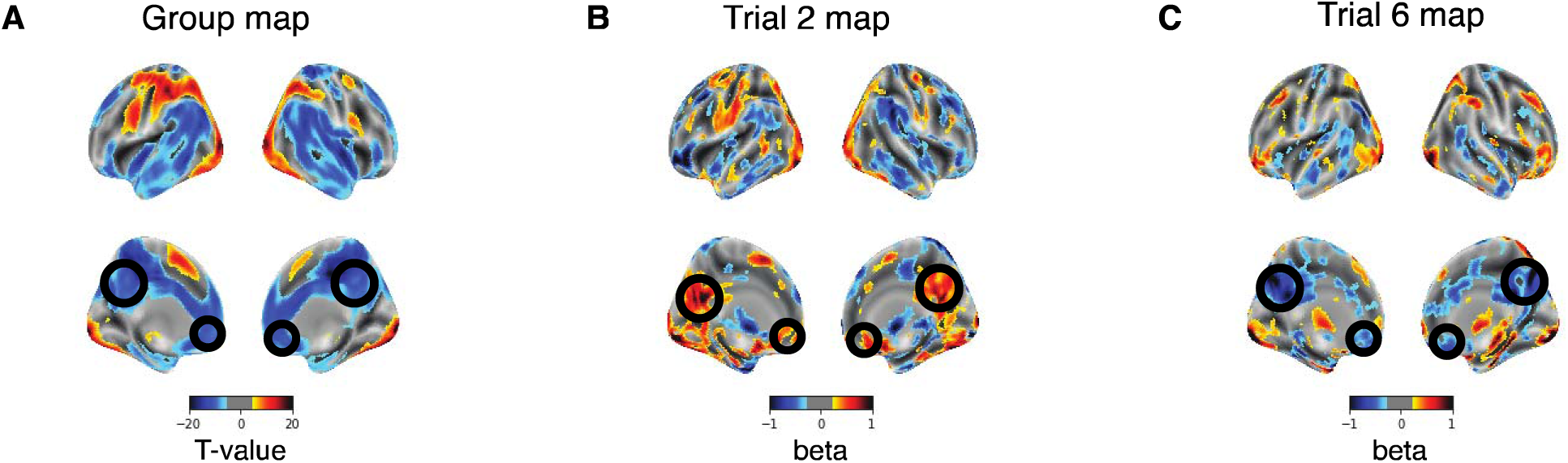
Variation in brain activity across individual trials in Experiment 1. (A) In Experiment 1, standard group analyses identified voxels with strong increases and decreases in task-induced brain activation across the cortex. The group brain map is thresholded at P < 0.001 for display purposes. Brain activation maps for (B) trial 6 and (C) trial 2 from subject 1 demonstrate substantial variability across trials that is not represented in the standard group brain map. Brain maps for the individual trials are thresholded at |beta| > 0.25 for display purposes. Black circles highlight the activation in the posterior cingulate cortex and orbitofrontal cortex. Panels B and C are shown for illustrative purposes only; formal analyses of the different activation patterns are shown in Figures 2-8.

**Table 1.** Experiment details.

|  | Subjects | Total # of Trials |
| --- | --- | --- |
| Experiment 1 | 50 | 35,682 |
| Experiment 2 | 39 | 31,024 |
| Experiment 3 | 40 | 9,959 |

### Multiple distinct but stable subtypes of trials during perceptual decision making

While single-trial activations are likely to be noisy and difficult to interpret, the divergence in brain activity between the two trials and to the group map highlights the possibility that more than one pattern of brain activation may exist when performing a task. To test for this possibility, we utilized a data-driven classification method to determine if multiple unique patterns of activation emerge across trials. For each trial, we estimated the task-induced brain activity in each voxel and pooled trials across subjects. We estimated the similarity across the activations between pairs of trials using Pearson correlation and clustered all trials using modularity-maximization to identify consistent activation patterns^22^. Clustering produced three subtypes of trials in Experiment 1 (**Fig. 2A)**. Each subtype accounted for a roughly similar proportion of all trials and each subtype was present in all subjects and in each of the four stimulus conditions in that experiment (see Methods; **Fig. 2B**).

**Figure 2.**
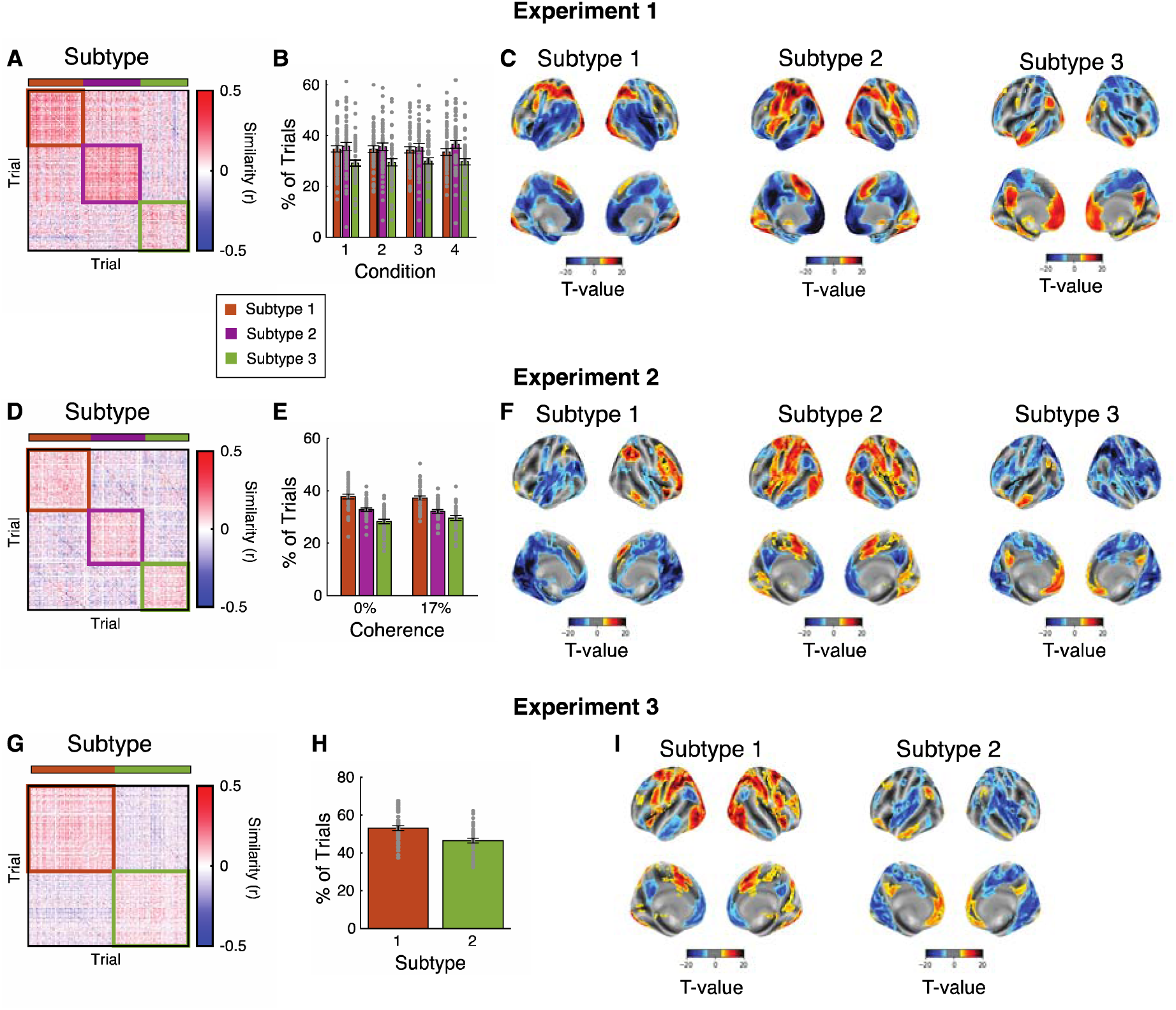
Subtypes of trials and activation maps in three perceptual decision-making tasks. (A-C) Results for Experiment 1. (A) Modularity-maximization based clustering identified three subtypes of trials. The colored squares correspond to the trials composing each subtype. Pearson correlation was used to calculate the spatial similarity in activation (betas) among individual trials. (B) The percent of trials classified as Subtype 1, 2, and 3 for each of the four stimulus conditions. The dots represent individual subjects. (C) Activation maps for each subtype. Each activation map was calculated by first averaging the trials for each subtype within a subject, followed by one-sample t-test to identify regions in which brain activity increased or decreased in response to the task. Brain maps are thresholded at P_FDR-corrected_ < 0.01. (D-F) Results for Experiment 2. (D) Modularity-maximization clustering identified three subtypes of trials. (E) The percent of trials classified as Subtype 1, 2, and 3. (F) Activation maps for each subtype. (G-I) Results for Experiment 3. (G) Modularity-maximization clustering identified two subtypes of trials. (H) The percent of trials classified as Subtype 1 and 2. Note that Experiment 3 contained only a single condition. (I) Activation maps for each subtype.

Critically, we examined the pattern of activation present in each subtype. To do so, we averaged the trials from each subtype within a subject. We then performed a group-level one-sample t-test on the average beta values to identify voxels with significant positive or negative activations (using a threshold of FDR-corrected P < 0.01; **Fig. 2C**). In Experiment 1, we found that Subtype 1 was characterized by bilateral visual, parietal and left motor activations, as well as medial frontal, cingulate, temporal, and right motor deactivations. Subtype 2 had many similarities with Subtype 1 – such as visual and parietal activations couple with medial frontal, cingulate, and anterior temporal deactivations – but featured strong bilateral activations in the insula (whereas Subtype 1 had bilateral deactivations in the insula). Subtype 3 had the most surprising profile with strong activations along the default mode network (DMN) and deactivations in a number of task-positive parietal and frontal areas. To further confirm these results, we performed a standard GLM analysis with subtypes as factors in the regression and found similar patterns of activations (**Fig. S1A**). We also confirmed these results using a standard cluster-based correction instead of a voxelwise FDR correction (**Fig. S1B**). Further, we performed the analysis within each of the four stimulus conditions separately to ensure that the subtypes do not simply reflect differences between conditions. We again found multiple activation patterns for each condition separately, with one of the patterns exhibiting strong DMN activation (**Fig. S2**).

To corroborate these findings, we performed the same analysis in two additional experiments that involved perceptual decision-making tasks – Experiments 2 and 3 (**Fig. 2D-I**). In Experiment 2, the group-level one-sample t-test on the average beta values identified voxels with significant positive and negative activations (FDR-corrected P < 0.01; **Fig. 2F**). Similar to Experiment 1, Subtypes 1 and 2 were characterized by activation across task-positive regions, whereas Subtype 3 exhibited strong activations in the DMN. In Experiment 3, clustering identified two subtypes. Similar to Experiments 1 and 2, the Subtype 1 exhibited activation in task-positive regions, whereas Subtype 2 exhibited strong activations in DMN (FDR-corrected P < 0.01; **Fig. 2I**). Taken together, these results confirm the existence of multiple activation patterns in three different experiments. Critically, in every experiment, one of the patterns exhibits activation in areas associated with the DMN, brain regions commonly thought to deactivate during tasks that require externally focused attention.

One possible reason for observing different patterns of activations is individual differences. That is, it is possible that each subject only exhibits a single pattern of activations, due to inter-subject differences, we find several different patterns within each experiment. However, the data do not support this possibility. As can be seen in Fig.2B,E,H, every subject has a substantial proportion of trials from each subtype. Focusing specifically on the DMN-associated subtype, we find that it reflects on average 29.6 ± 8.9 (mean ± SD) of trials in Experiment 1, 35.8 ± 5.2% of trials in Experiment 2, and 46.5 ± 8.1% of trials in Experiment 3. The relatively small SD values demonstrate that all subjects have a large proportion of trials reflecting the DMN-associated subtype. In other words, while there are likely meaningful individual differences in how much each subtype is represented, the existence of different subtypes is not in itself a function of individual differences between subjects.

To further understand the nature of each subtype, we examined the pattern of activation within established brain networks for each subtype. For each subject we averaged the voxels within each of the seven brain networks – frontoparietal network (FPN), default mode network (DMN), dorsal attention network (DAN), limbic network (LIM), ventral attention network (VAN), somatomotor network (SOM), and visual network (VIS) – defined in the Schaefer Atlas^23^. We found that Subtype 3 exhibited the strongest activation in the DMN and LIM networks but weakest activations in the FPN, DAN, and VIS networks (one-samples t-tests; P_FDR-corrected_; **Fig. 3A**). Subtypes 1 and 2 both showed strong activation in the DAN and VIS networks, but differed in the activation strength across the FPN, VAN, and SOM networks. We performed the same analysis in Experiments 2 and 3, and found results broadly similar to Experiment 1 (one-sample t-test; P_FDR-corrected_ < 0.01; **Fig. 3B,C**).

**Figure 3.**
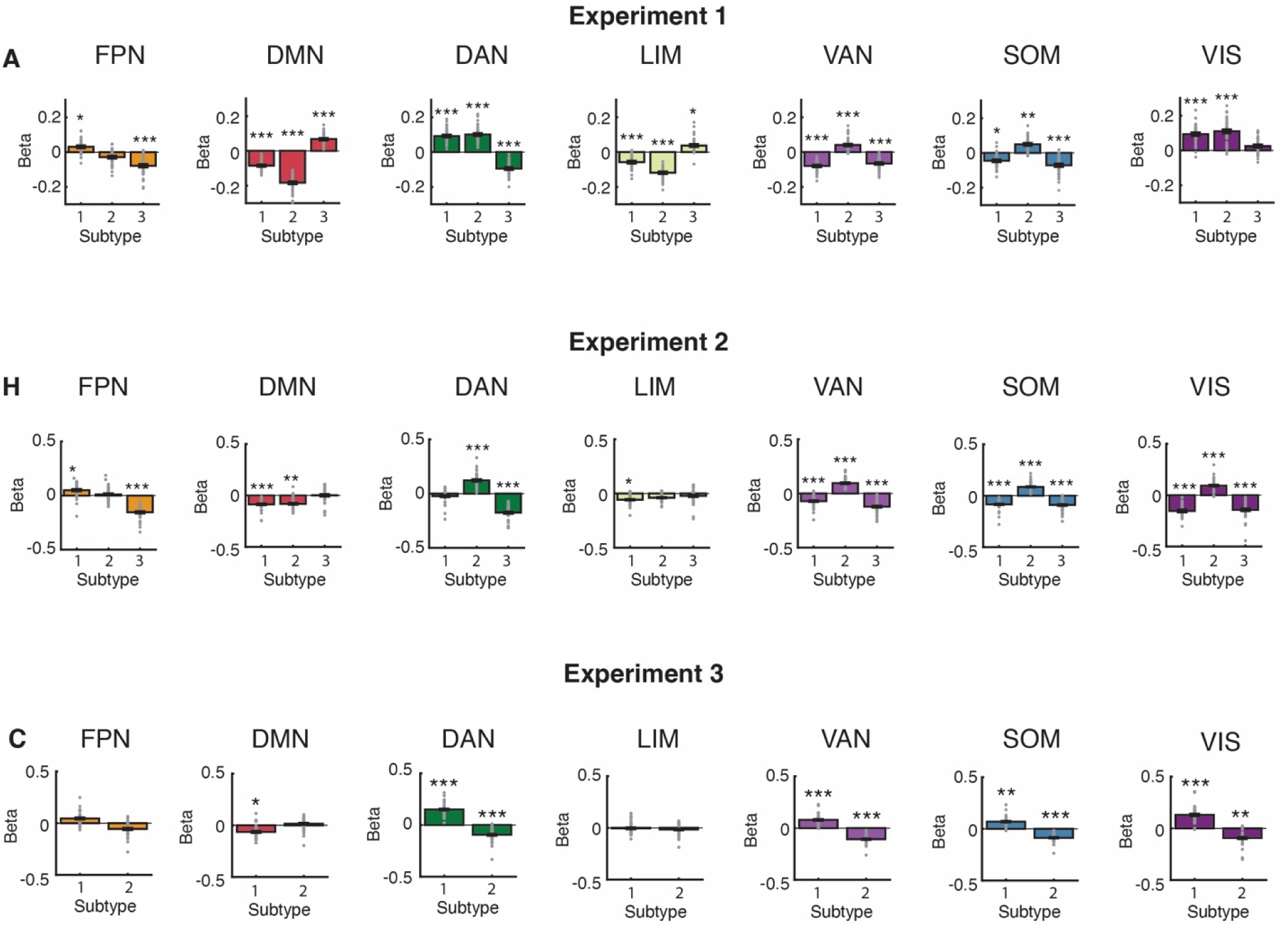
Activation for each of seven large-scale brain networks for each subtype. (A) Results for Experiment 1. Activation for each of seven large-scale brain networks for each subtype. The bar graph shows the average change in activation (mean ± sem). The grey dots show the activation in each subject. (B) Results for Experiment 2. (C) Results for Experiment 3. Activation changes from baseline were estimated using a one-samples t-test. *** P_FDR-corrected_ < 0.001; ** P_FDR-corrected_ < 0.01; * P_FDR-corrected_ < 0.05. FPN, Frontal Parietal Network; DMN, Default Mode Network; DAN, Dorsal Attention Network; LIM, Limbic Network; VAN, Ventral Attention Network; SOM, Somatomotor Network; VIS, Visual Network.

### Subtypes are robust to methodological choices, noise, and experimental factors

Importantly, we confirmed that the multiple activation patterns identified in each of the different tasks could not be explained by methodological choices, noise in the data, or experimental factors.

First, we checked whether the obtained clusters depend on methodological choices related to the clustering algorithm we used. Specifically, the number of clusters identified using modularity-maximization depends on the value of the resolution parameter (γ), which was set to its standard value of 1 in the above analyses^24^. To determine that the obtained clusters are robust to this value, we re-ran the analysis using a range of gamma values from 0.8 to 1.2 to determine if this parameter affects the composition of the clusters. We found the number of clusters was stable for gamma values in the range of 0.8 to 1.01 for all three experiments (**Fig. 4A**). Gamma values higher than 1.01 led to more clusters but without affecting the core subtypes. Instead, these high gamma values simply led to a small number of trials from each subtype to be separated into new clusters (**Fig. 4B**). These results demonstrate that the existence of the core clusters does not depend on the value of the resolution parameter (γ).

**Figure 4.**
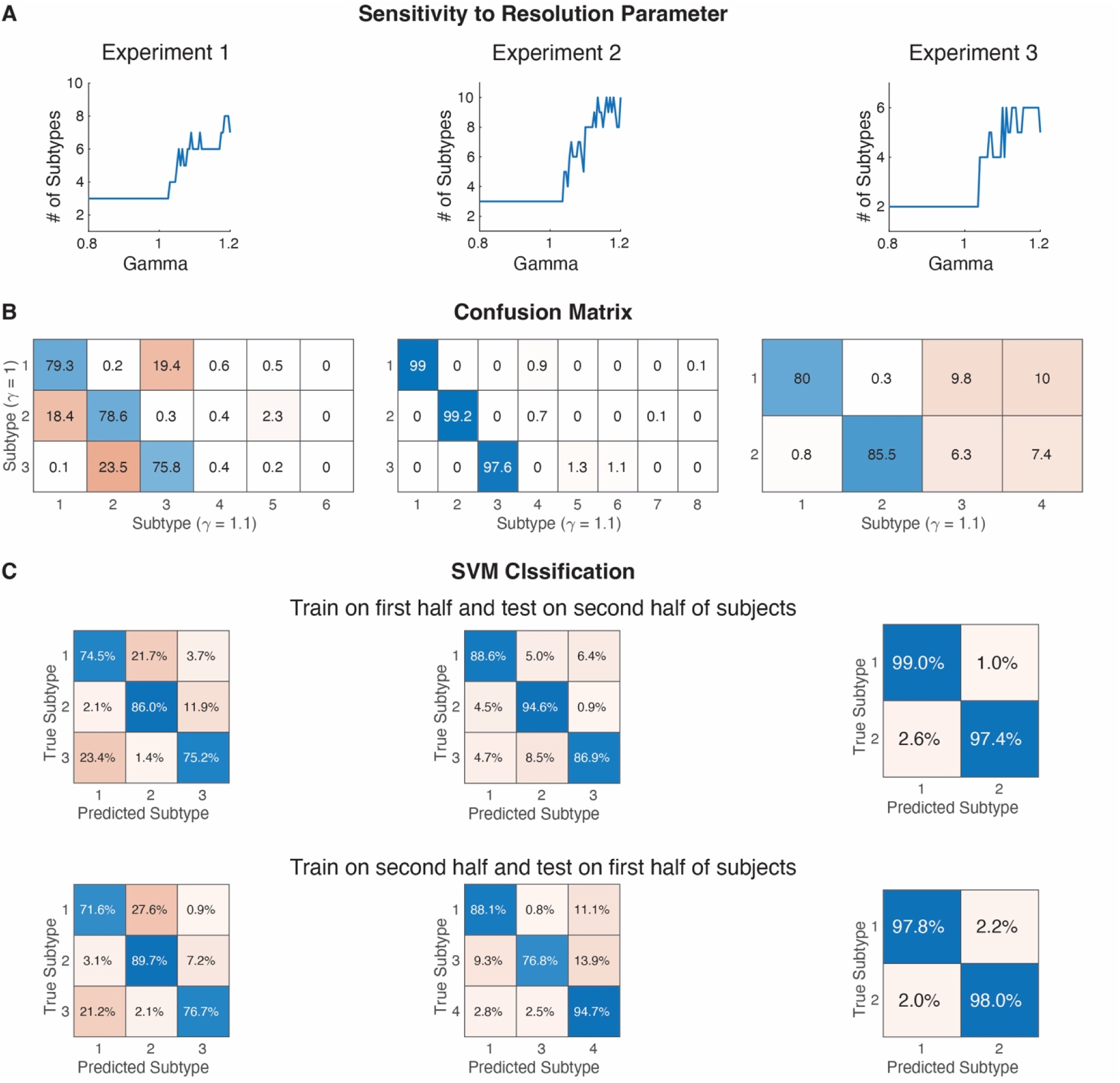
Subtypes are independent of methodological choices. (A) Sensitivity of clustering to resolution parameter. The number of clusters (subtypes) was stable over a range of resolution parameter (γ) values from 0.8 to 1.01 for each experiment. Higher γ values resulted in a larger number of clusters. (B) The increased number of subtypes obtained with higher γ levels arise from separating a few trials from the main subtypes. To demonstrate that, we compared the clusters obtained with γ = 1 and γ = 1.1. As the figure demonstrates, there is a strong mapping between the first three (Experiments 1 and 2) or first two (Experiment 3) subtypes obtained with γ = 1 and γ = 1.1. Thus, higher γ values do not lead to qualitatively different subtypes. (C) SVM Classification. The SVM classifier correctly labeled on average 87.2% of trials across all tasks.

Second, we confirmed that our results do not stem from noise in the data. To do so, we performed a range of analyses. We began by confirming that the clustering remains stable when performed on different subjects. Specifically, we split the subjects in half and then repeated the clustering analysis on each half. We then trained an SVM classifier to predict the labels on the data from one half of the subjects using the labels for the other half. The SVM classifier correctly labeled on average 87.2% of trials across Experiment 1-3 (**Fig. 4C**). By comparison, an SVM classifier trained to separate trials based on experimental condition performed at chance (**Fig. S3**). Importantly, we also confirmed that the clusters were not related to head motion. Specifically, we found no clear relationship between subtypes and fMRI noise as measured with Frame Displacement, temporal derivative of the time course (DVARS), or each of the six motion parameters. To be as sensitive as possible, we conducted a series of pairwise comparisons between every two subtypes in every measure of subject motion. Focusing on Experiment 1, we observed no significant differences between each of the motion estimated parameters among the subtypes (**Fig. 5**). Extending this analysis to Experiment 2 and 3, when using uncorrected tests, we obtained two significant differences from a total of 126 tests run across all three experiments, which is less than the 5% expected rate of significant results assuming no true effects (see Supplemental Results and **Fig. S4-S5**). Finally, because voxel-wise estimates can be unstable and noisy, we repeated the clustering analyses using average activations within 200 brain regions and still found similar results (see Supplemental Results and **Fig. S6**).

**Figure 5.**
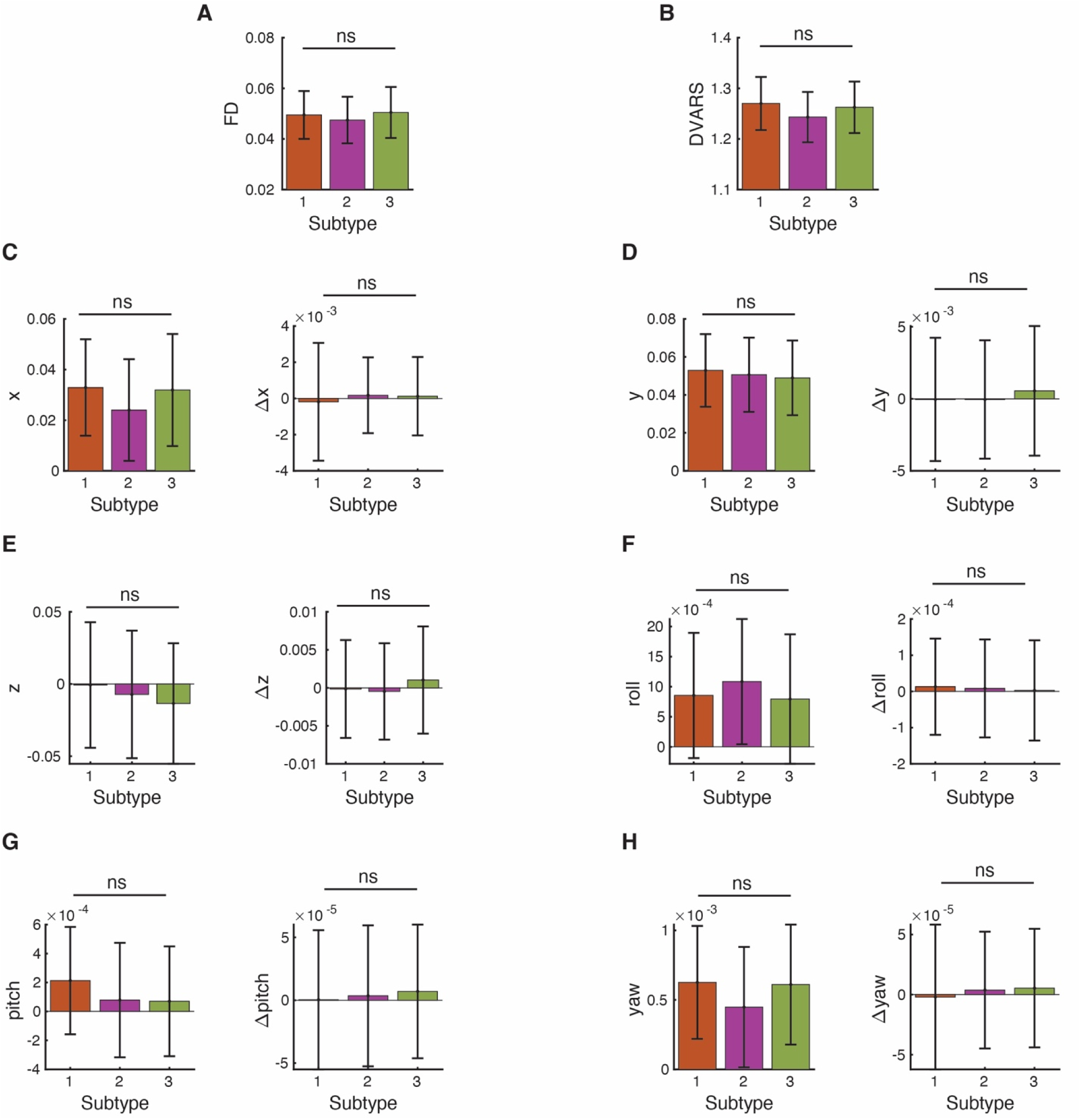
No differences in head motion between subtypes in Experiment 1. The average (A) Frame displacement (FD), (B) DVARS, (C) x-, (D) y-, (E) z-, (F) roll-, (G) pitch-, (H) yaw-direction per subtype. For each trial we estimated 14 different motion-associated artifacts. Estimated motion values were averaged per subtype within a subject and statistical differences were determined using paired-samples t-test without any correction. For panels C-H, right panels show the 1^st^ derivatives. ns, not significant.

Third, we confirmed that experimental factors including subject age and sex, trial position, or the time interval between successive trials were not driving the activation difference between trials (see Supplemental Results and **Fig. S7-S8**). Collectively, we found that none of these factors plays a substantial role in determining the obtained subtypes. Taken together, these results suggest that the multiple activation patterns are not simply driven by trivial experimental factors, subject characteristics to various types of noise or experimental factors that might affect trial-by-trial activation patterns over the course of a task.

### Behavioral differences between subtypes

Having identified these three different subtypes of trials in Experiment 1, we investigated how they affected behavioral performance. We compared how behavioral performance differed among subtypes using a mixed-effect model to account for the different conditions within the experiments. The model assessed the effects of subtype and condition with the subject as a random factor on behavioral performance. For Experiment 1, significant effects of subtype were present for reaction time (RT) (t(35679) = 2.90, P = 0.004) and confidence (t(35679) = -3.37, P = 0.001), but not accuracy (t(35679) = -1.78, P = 0.07).

The same analyses for Experiment 2 uncovered significant differences in RT (t(30351) = -4.99, P = 5.83 x 10^-7^) and confidence (t(30351) = 4.37, P = 1.25 x 10^-5^), but not accuracy (t(30351) = 0.33, P = 0.74). Interestingly, and in contrast to Experiment 1, in Experiment 2, the fastest RTs were associated with Subtype 3, which showed activation in the DMN. Lastly, in Experiment 3, there were no significant difference in accuracy or RT between subtypes (P > 0.05). (Note that confidence was not measured in the Experiment 3.) Follow-up pairwise comparison analyses using paired t-test reflected the results obtained with the mixed-effect model (**Fig. 6**). Overall, the differences in accuracy, RT, and confidence were small and their direction was sometimes inconsistent between studies, suggesting that the subtypes do not simply reflect differences in task difficulty or other stimulus characteristics.

**Figure 6.**
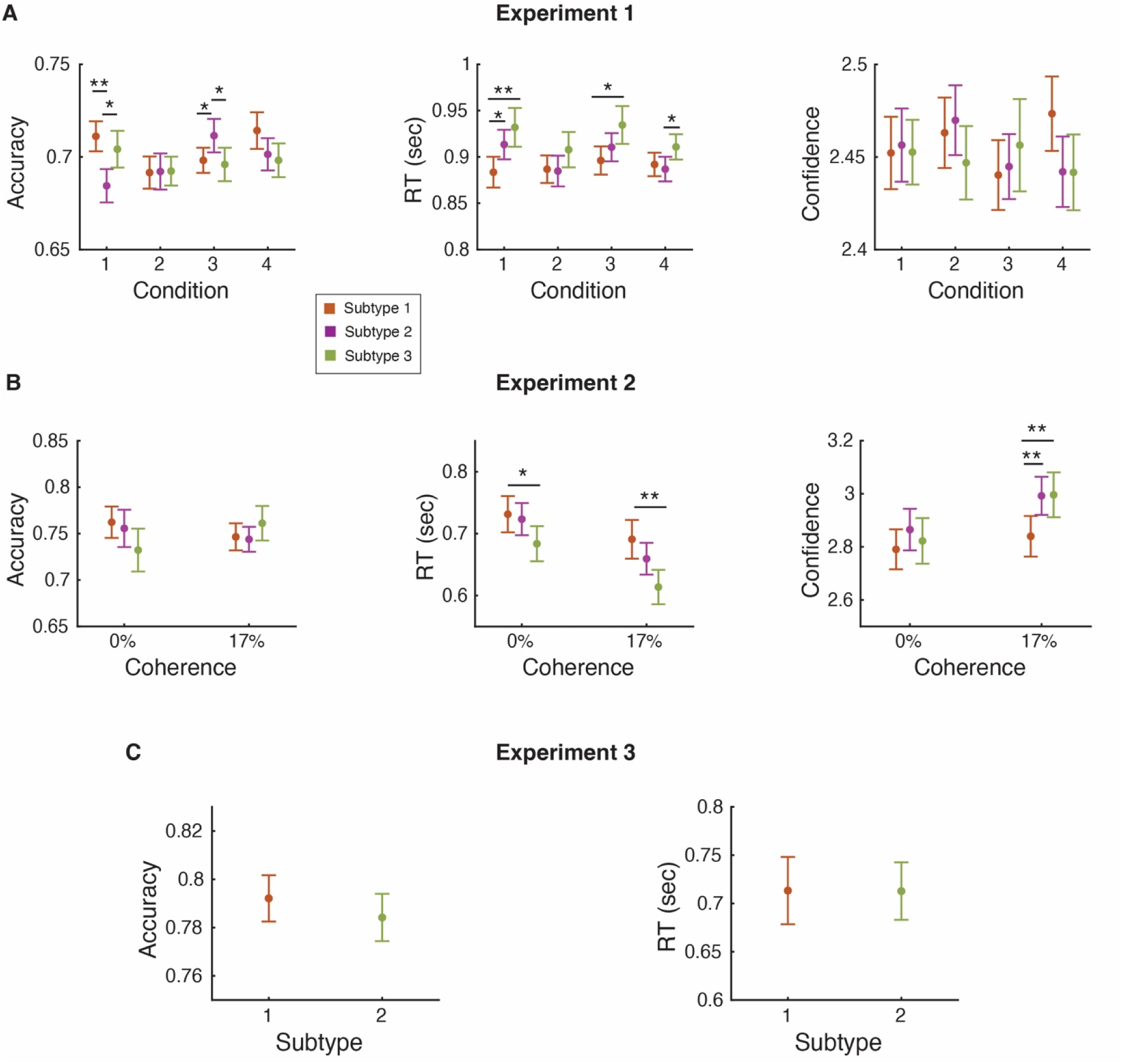
Behavioral differences between subtypes. Differences between subtypes in accuracy, RT, and confidence for (A) Experiment 1, (B) Experiment 2, and (C) Experiment 3. Statistical testing was conducted using linear mixed-effect models where the effects of subtype and condition were fixed effects and subject was a random factor. For post-hoc analysis, first behavioral measures were averaged within a subject and a paired-samples t-test was used to determine significant differences. Averaging within a subject result in loss of power compared the mixed-effected model which was conducted at the trial level. Note that Experiment 3 contained only a single condition and confidence was not measured. Error bars show SEM. ** P < 0.01; * P < 0.05.

Further, we tested if clustering trials improved the brain-behavior relationship estimation. Focusing on Experiment 1 since it contained the most trials, we estimated the change in sensitivity in the brain-behavior relationship within each brain region part of the Schaefer 200 brain atlas. Specifically, within each subject, for each region we correlated the trial activation strength with accuracy, RT and confidence for each subtype of trials and all trials together. We found that the correlation between behavioral measures (accuracy, RT, and confidence) and brain activation improved (P < 0.05) for each subtype compared to when all trials were considered together, suggesting that considering each subtype separately increases the sensitivity in brain-behavior relationship (**Fig. 7**).

**Figure 7.**
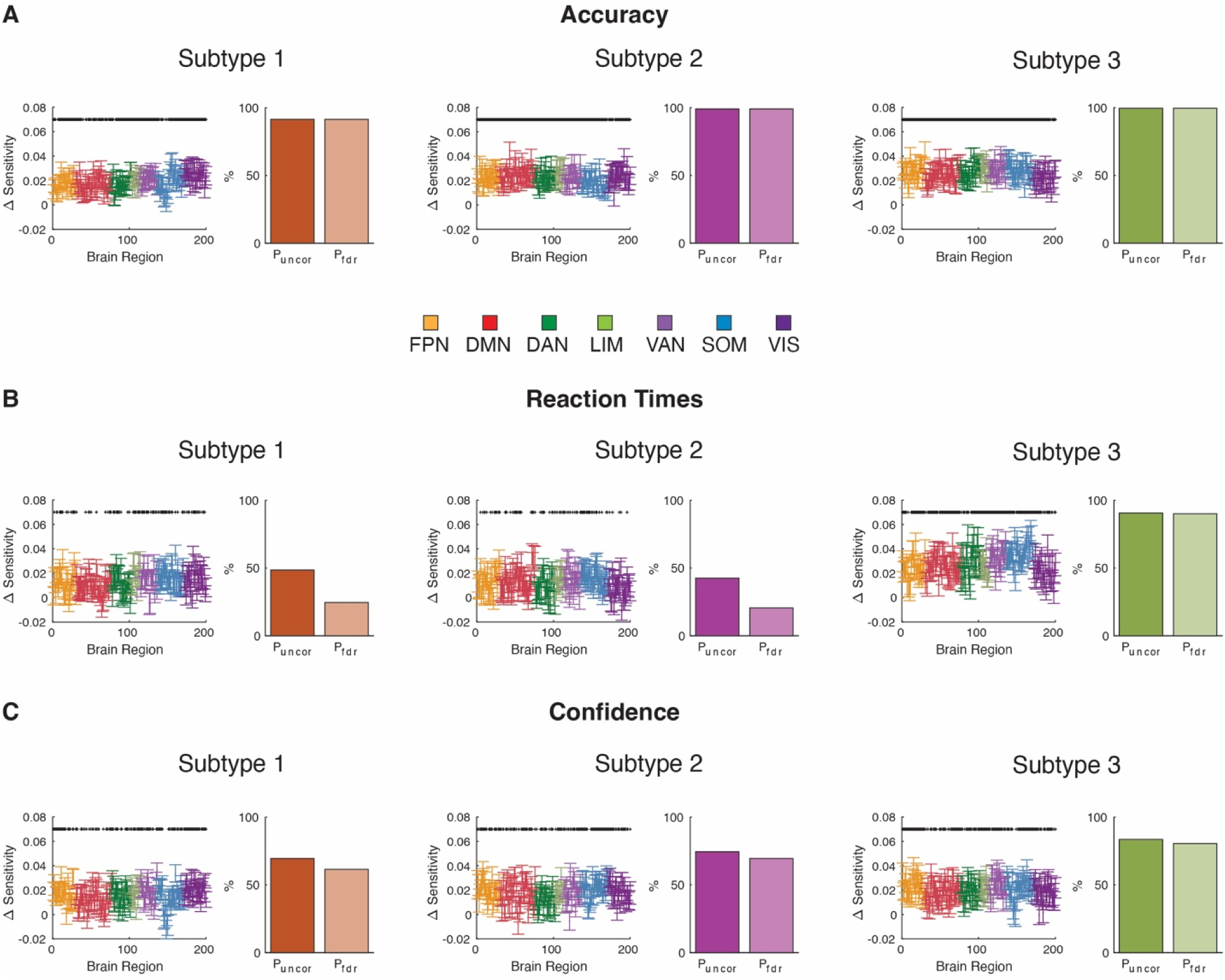
Subtyping trials improves sensitivity in the relationship between accuracy, RT, and confidence with activation in Experiment 1. The correlation difference between average activation strength in a brain region part of the Schaefer 200 atlas and (A) accuracy, (B) RT, and (C) confidence for each subtype compared to all trials together. Significant differences were estimated with group level one-sample t-tests. The dot plots on the left show the average increase in sensitivity in the brain-behavior correlation across subjects (mean ± sem). The bar plots on right show the percentage of regions that the sensitivity significantly increased without any correction for multiple comparisons (P_uncor_) and with a false discovery rate multiple comparison correction (P_FDR_). *, P_uncor_ < 0.05.

### Processes underlying subtype activation

Having established the existence of subtypes that differ in their neural activation patterns and behavioral correlates, we examined the transitions between different subtypes and potential mechanisms that can lead to the emergence of these subtypes. We first investigated the transition probabilities between subtypes to understand whether the different subtypes were randomly intermixed or whether their occurrence followed a specific pattern. We found that a trial of a specific subtype was much more likely to be followed by a trial of the same subtype (**Fig. S13**). These results suggest that the subtypes reflect slow changes in underlying cognitive processes. Additionally, we build a model that can generate the multiple activation patterns from the structural or functional connectivity, both of which have been shown to be capable of predicting brain activation^25–27^. The result from the model suggest that global brain activation is primarily driven by the stimulus-drive from a few networks (see Supplementary Methods, Supplementary Results, and **Fig. S14-S15**).

### Brain regions exhibiting consistent activation across trials

Besides exploring the differences between the trials, we also examined what is common across them. To identify areas exhibiting consistent activations or deactivations in brain activity across trials, we identified the voxels in which the sign of activation was always in the same direction. Specifically, we first binarized the activation the brain maps patters for each trial and identified the voxels for which the sign of activation always in the same direction for all trials in a given subject. We then plotted voxels that have the same activation sign in all trials in a large proportion of all subjects. As may be expected, we found consistent activations in the visual and left motor cortex, as well as consistent deactivations in medial somatomotor, right motor, and bilateral temporal cortex (**Fig. 8**). These results confirm that despite the existence of different subtypes, expected activation effects remain robust across individual trials.

**Figure 8.**
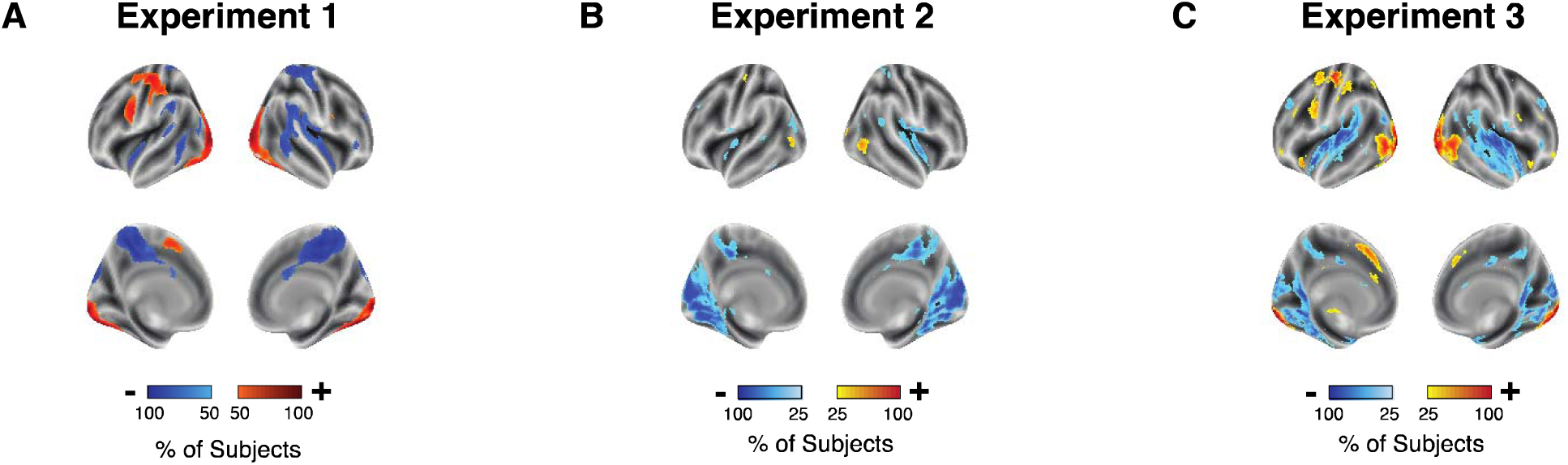
Maps of activations that are consistent across trials. Voxels exhibiting consistent activation across trials in (A) Experiment 1, (B) Experiment 2, and (C) Experiment 3. We first binarized the activation in the brain maps patters for each trial and identified the voxels for which the sign of activation always in the same direction for a given subject. We then plotted voxels that have the same activation sign in all trials in a large proportion of all subjects. For visualization purposes, maps are threshold at 50% of subjects in Experiment 1 and 25% of subjects in Experiments 2 and 3.

### Extension to a working memory task

Finally, beyond the three perceptual decision-making experiments, we also examined if multiple activation patterns exist in a different cognitive task. Specifically, we analyzed the *n*-back task data from the Human Connectome Project where subjects completed equal number of 0- and 2-back trials^28^. In the same manner as for Experiments 1-3, for each trial we estimated the task-induced brain activity in each voxel and pooled trials across subjects and *n*-back conditions, and then clustered the trials using modularity-maximization. Clustering identified three subtypes present in both the 0- and 2-back conditions. However, two of these clusters were relatively similar to each other, while the third one appeared in few trials and may reflect motion artifacts (see Supplementary Methods and Results for details **Fig. S9-S12**). These results suggest that very different patterns of activity may not always emerge in all tasks.

## Discussion

Fluctuations in brain activity are ubiquitous during simple and cognitively demanding tasks^16,29^ and are often indicative of variation in cognitive processing^30^. However, meaningful variation in internal states that impacts cognition and behavior remains challenging to discover and characterize. Here we examined if distinct patterns of brain activity emerge on different trials when accomplishing the same task. We utilized a data-driven clustering method based on modularity-maximation to identify consistent patterns of brain activity on individual trials. Across three perceptual decision-making experiments, the clustering analysis identified multiple discrete subtypes of trials. Surprisingly, in each of the three experiments, one of the subtypes exhibited activations in DMN. To the best of our knowledge, this is the first analysis to report DMN activation on a subset of trials during a task that requires *externally focused* attention. These findings demonstrate that the same task can be accomplished in the presence of widely varying brain activation patterns.

Our most striking finding was that trials from one of the subtypes in Experiments 1-3 showed a strong increase in DMN activity even though subjects were engaged in a perceptual decision-making task that canonically leads to DMN suppression. Behaviorally, the DMN-associated subtype was characterized by accuracy, RT, and confidence that were similar to the other subtypes associated with task-positive regions. These behavioral results are particularly surprising because DMN activation is usually associated with mind-wandering and being off-task^31,32^. It is important to note that DMN is known to be activated during task requiring internal or self-relevant focus^33–37^ or boredom^38^. As a result, the activation of the DMN may reflect a change in attention from externally to internally focused attention or boredom. Yet such interpretations would presumably predict larger behavioral differences between the DMN-associated subtype and the subtypes associated with increased activation in task-positive regions. Our findings demonstrate substantial DMN activations even for tasks that canonically should deactivate the DMN, which challenges the notion that DMN should be viewed as a task-negative network^39^.

Why are multiple qualitatively different patterns of brain activity emerging from different trials of the same task? One may speculate that a cognitive process can be accomplished through multiple pathways. In fact, it has been hypothesized that multiple cognitive pathways could exist because of degeneracy^19^, and this hypothesis is supported by work utilizing theoretical models and patient populations^20^. Similar to a person driving home from work, there may be multiple routes to get home, and the path taken may depend on various factors such as the extent of traffic on a specific route. In this interpretation, the subtypes in perceptual decision-making tasks may be characterized as reflecting decision-making pathways in the presence of endogenous attention^40–42^, exogenous attention^43,44^, or internally focused attention^37^.

Instead of discrete pathways, another possibility is that the trial subtypes are part of a gradient in activation across the brain^45,46^. In this interpretation, the trial subtypes would indicate the axes of this space. Specifically, in the perceptual decision-making tasks, the location of an individual trial within this space could be dependent on the extent of top-down control^47,48^, the salience of the stimulus^42^, and internally oriented attention^49,50^. Critically, neither interpretation excludes the existence of shared components. Indeed, we found that the visual regions exhibited similar patterns in activation in all subtypes across all experiments. Additionally, this consistent pattern of activation included regions in the frontal, parietal, and temporal areas, suggesting a core component across all trials. Future research should examine the underlying processes that lead to the emergence of these trial subtypes across an array of cognitive functions.

The current work has several limitations. First, it is currently unclear why the subtypes across the three experiments showed some important differences. These differences could reflect subtle differences in the stimuli (e.g., color stimuli in Experiment 1 vs. motion stimuli in Experiments 2 and 3), task (e.g., subjects rated confidence in all runs of Experiment 1, in half the runs of Experiment 2, and did not rate confidence at all in Experiment 3), or other aspects of the experimental design. Future studies can examine how the subtypes change in response to specific experimental factors. Second, while we suspect that the subtypes reflect different cognitive processes, it is currently difficult to identify what specifically these processes are. Behaviorally, we could only examine accuracy, RT, and confidence, generally finding only subtle differences among the subtypes. It is likely that the subtypes reflect other signatures of internal processing such as pupil dilation, skin conductance, or heart rate variability, but data on these variables was not available in the current datasets. Future research should examine a wider range of behavioral and physiological differences between trial subtypes.

In conclusion, we found that both perceptual decision-making and working memory tasks featured multiple distinct patterns of brain activation. These results suggest that several independent pathways may be employed to accomplish a task.

## Methods

### Experiment 1 Subjects and Task

Fifty healthy subjects (25 females; Mean age = 26; Age range = 19-40) that has been described elsewhere^51^. All subjects were screened for any history of neurological disorders or MRI contraindications. The study was approved by Ulsan National Institute of Science and Technology Review Board and all subjects gave written consent.

Subjects had to determine which set of colored dots (red vs. blue) is more frequent in a cloud of dots. Each trial began with a white fixation dot presented for a variable amount of time between 500-1500 ms. Then, the stimulus was shown for 500 ms, followed by untimed decision and confidence screens. The stimulus consisted of between 140 and 190 red- and blue-colored dots (dot size = 5 pixels) dispersed randomly inside an imaginary circle with a radius of 3° from the center of the screen. Four different dot ratios were used – 80/60, 80/70, 100/80, and 100/90, where the two numbers indicate the number of dots from each color. The experiment was organized in blocks of 8 trials each, with each dot ratio presented twice in a random order within a block. The more frequent color was pseudo randomized so that there were equal number of trials within a block where red or blue was the correct answer. The luminance between the red and blue dots was not matched. Subjects performed a total of 768 trials. Three subjects completed only half of the 6^th^ run and another three subjects completed only the first 5 runs due to time constraints. The remaining 44 subjects completed the full 6 runs.

### Experiment 1 MRI Recording

The MRI data was collected on a 64-channel head coil 3T MRI system (Magnetom Prisma; Siemens). Whole-brain functional data were acquired using a T2*-weighted multi-band accelerated imaging (FoV = 200 mm; TR = 2000 ms; TE = 35 ms; multiband acceleration factor = 3; in-plane acceleration factor = 2; 72 interleaved slices; flip angle = 90°; voxel size = 2.0 x 2.0 x 2.0 mm^3^). High-resolution anatomical MP-RAGE data were acquired using T1-weighted imaging (FoV = 256 mm; TR = 2300 ms; TE = 2.28 ms; 192 slices; flip angle = 8°; voxel size = 1.0 x 1.0 x 1.0 mm^3^).

### Experiment 2 Subjects and Task

Thirty-nine subjects (23 females, average age = 21.5 years, range = 18–28 years, compensated $50 for participation) were instructed to indicate whether a moving-dots stimulus had an overall coherent motion (always in downward direction) or not. Subjects had no history of neurological disorders and had normal or corrected-to-normal vision. The study was approved by the Georgia Tech Institutional Review Board. All subjects were screened for MRI safety and provided informed consent. The study’s method and procedure were carried out according to the declaration of Helsinki.

Detailed description can be found in Yeon *et. al*.,^52^. In brief, each trial began with a fixation mark presented randomly for 1, 2, or 3 sec and followed by the stimulus presented for 500 ms. In the first half of the experiment (runs 1–3), subjects performed the task and were never told to evaluate their confidence level. In the second half of the experiment (runs 4–6), subjects made their perceptual decision and immediately after were asked to indicate their confidence level.

### Experiment 2 MRI Recording

The MRI data were collected on 3T Prisma-Fit MRI system (Siemens) using a 32-channel head coil. Anatomical images were acquired using T1-weighted sequences (MEMPRAGE sequence, FoV = 256 mm; TR = 2530 ms; TE = 1.69 ms; 176 slices; flip angle = 7°; voxel size = 1.0 × 1.0 × 1.0 mm^3^). Functional images were acquired using T2*-weighted gradient echo-planar imaging sequences (FoV = 220 mm; TR = 1200 ms; TE = 30 ms; 51 slices; flip angle = 65°; voxel size = 2.5 × 2.5 × 2.5 mm^3^).

### Experiment 3 Subjects and Task

The analysis was based on 40 subjects who performed a motion discrimination task. Subjects were compensated $20/hour or 1 course credit/hour for a total of 2.5 hours. All subjects were right-handed with normal hearing, normal or corrected-to-normal vision, had no history of neurological disorders, brain trauma, psychiatric illness, or illicit drug use. The study was approved by the Georgia Tech Institutional Review Board. All subjects were screened for MRI safety and provided written informed consent.

Detailed description can be found in Haddara & Rahnev^53^. In brief, subjects judged the motion direction (left or right) of white dots (density: 2 dots/degree², speed: 5°/s) presented in a black circle (3° radius) in front of a grey background. A proportion of dots moved coherently in the right or left direction while the rest of the dots moved randomly. Each dot motion stimulus was preceded by a letter cue (“L” = Left, “R” = Right, “N” = Neutral). The letters L and R predicted the forthcoming stimulus with 75% validity, whereas the letter N was not predictive (both left and right motion were equally likely). Each trial began with cue presentation for 2, 4, or 6 seconds (chosen randomly), followed by a 3-second dot motion stimulus and an untimed response. A screen with a fixation dot was then presented between trials for a period of 1 or 2 seconds.

### Experiment 3 MRI Recording

BOLD fMRI signal data was collected on a 3T MRI system (Prisma-Fit MRI system; Siemens) using a 32-channel head coil. Anatomical images were acquired using T1-weighted sequences (MEMPRAGE sequence, FoV = 256 mm; TR = 2530 ms; TE = 1.69 ms; 176 slices; flip angle = 7°; voxel size = 1.0 x 1.0 x 1.0). Functional images were acquired using T2*-weighted gradient echo-planar imaging sequences (FoV = 220 mm; slice thickness = 2.5, TR = 1200 ms; TE = 30 ms; 51 slices; flip angle = 65°; voxel size = 2.5 x 2.5 x 2.5, multi band factor = 3, interleaved slices).

### Experiment 1-3 MRI Preprocessing

MRI data were preprocessed with SPM12 (Wellcome Department of Imaging Neuroscience, London, UK). We first converted the images from DICOM to NIFTI and removed the first three volumes to allow for scanner equilibration. Following standard practice, we preprocessed with the following steps: de-spiking, slice-timing correction, realignment, segmentation, coregistration, normalization, and spatial smoothing with 10 mm full width half maximum (FWHM) Gaussian kernel except for Experiment 3 where smoothing was performed with a 6 mm FWHM Gaussian kernel. Despiking was done using the 3dDespike function in AFNI. The preprocessing of the T1-weighted structural images involved skull-removal, normalization into MNI anatomical standard space, and segmentation into gray matter, white matter, and cerebral spinal fluid, soft tissues, and air and background. The segmentation of T1-weighted images was conducted in SPM12 with default parameters and normalized to the default MNI template with 2 cm^3^ voxel dimensions.

### Single-Trial Beta Estimation

Single-trial beta responses were estimated with a general linear model (GLM) using GLMsingle, a Matlab toolbox for single-trial analyses^21^. The hemodynamic response function was estimated for each voxel and nuisance regressors were derived in the same manner as previously described in Allen *et. al.,* ^54^. Additionally, regressors for the global signal and for six motion parameters (three translation and three rotation) were included. The single-trial betas were estimated in three batches. In each batch, the betas for every third trial were estimated because the trials in our study were temporally close together. Also, trials that were within 20 seconds from the end of run were removed. The betas for each voxel represent the estimated trial-wise BOLD response and are relative to the BOLD signal observed during the absence of the stimulus^21^.

### Modularity-maximization Based Clustering

The beta maps per trials were pooled among subjects to ensure that there was consistency in clustering correspondence. A trial-by-trial similarity matrix was created using the Pearson Product Correlation using all gray-matter voxels, except in the working memory task where all brain voxels were used. Clustering of the similarity matrix was conducted using modularity-maximization^22^. Modularity-maximization does not require the number of clusters to be specified and the resolution of the clusters was controlled with resolution parameter, γ = 1. Modularity-maximization was implemented with the Generalized Louvain algorithm part of the Brain Connectivity Toolbox^55^.

The community detection method used in the analysis is not deterministic and the results can depend on the specific random seeds. Crucially, as examined by Lancichinetti et al.^56^, these limitations can be overcome using consensus clustering to identify stable clusters out of a set of partitions. Moreover, in our previous work of clustering single trials, we had found that 100 iterations were sufficient to identify stable clusters^57^. Specifically, consensus clustering identifies a single representative partition from the set of 100 iterations. This process involves the creation of a thresholded nodal association matrix which describes how often nodes were placed in the same cluster. The representative partition is then obtained by using a Generalized Louvain algorithm to identify clusters within the thresholded nodal association matrix^58^. We have previously utilized this method to identify trial subtypes in brain activity measured with electroencephalography in a motion discrimination^59^ task and working memory task^57^.

To ensure that the number of clusters was not dependent on the value of the resolution parameter (gamma), we re-ran the clustering with gamma values ranging from 0.8 to 1.2. A value of 1 for the resolution parameter is considered standard^63^, which is why we used it. However, increasing gamma favors the identification of smaller clusters and lowering gamma favors the identification of larger clusters. Further, we expected to find a small number of clusters because in previous work in which we clustered trials in EEG data we found at most three^57,59^.

### Standard Group-Level Analysis

A standard task-based GLM analysis was conducted to identify voxels in which the beta values significantly deviated from zero. Specifically, a single activation brain map was created per subject by averaging the individual beta maps across trials and a one-sample t-tests was conducted across subjects to identify the regions that deviated from zero.

### Trial Subtype Activation

In a similar manner to the standard group-level analysis, a trial subtype task-based analysis was conducted to identify voxels in which the beta values for each subtype of trials significantly deviated from zero. Trials of the same subtype were averaged within each participant resulting in one average map per subtype for each participant. A group-level one-sample t-test was conduct for every voxel and p-values were FDR-corrected for multiple comparisons.

### Standard General Linear Modeling

To further corroborate the results, we also performed a standard GLM using SPM12 with subtypes as factors in the regression and found similar patterns of activations. We fit a GLM that allowed us to estimate the beta values for each voxel in the brain in each of the trial subtypes.

The model consisted of separate regressors for each of the subtypes, inter-block rest periods, as well as linear and squared regressors for six motion parameters (three translation and three rotation), five tissue-related regressors (gray matter, white matter, and cerebrospinal fluid), global signal, and a constant term per run.

### Determining the Effect of Preprocessing Choices on Results

Our analysis is based on re-analyzing existing data and pre-processing was done in the original analysis. The data we are re-analyzing had been smoothed with kernels between 6 and 10 mm full-width half-max (FWHM), which is standard practice for fMRI studies focusing on task-activation. In our main analyses, we used the previous smoothing values to avoid any flexibility in the analysis pipeline. To confirm that the smoothing level did not drive the results, we repeated the clustering analyses using a 4-mm FWHM smoothing kernel, which is less aggressive than in our main analysis. Additionally, we incorporated WM signal and CSF signal as nuisance regressors in this analysis to confirm that our results were not dependent on the WM and CSF. Yet, these control analyses found largely the same subtypes as in the main analyses (**Fig S1C**).

### Consistency in Activation Across Subtypes

To identify voxels exhibiting consistent task-induced changes in brain activity, we examined the consistency of the sign of voxel activations (positive or negative) across subjects. To do so, the brain maps of each trial were first removed all non-gray matter voxels. We then binarized the voxel activation values *actavatawn_i_* such that:

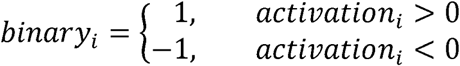

The consistency of the sign of a voxel’s activation across subtypes (*c_i_*) was then calculated as total number of trials for a which voxel *i* was positively or negatively activated using the formula:

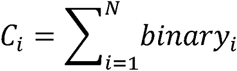

where *N* is the number of trials completed by a subject. Consequently, *c_i_* can take values between -*N* (all trials having negative activation for that voxel) to *N* (all trials having positive activations for that voxel). We then selected voxels where *c_i_*= ±*N* for which brain activity consistently increased or decreased across all trials. The brain maps where then averaged across subjects.

### Voxel and Brain Network Differences Between Subtypes

Differences in task-based brain activity between subtypes was conducted to identified voxels in which the beta values between subtypes differed. The analysis was conducted both at the voxelwise and between large-scale brain networks. For voxelwise analyses, a paired t-test was used to test for differences between subtypes. For the comparison between large-scale brain networks, the beta values from voxels associated with one of seven large-scale brain networks part of the Schaefer Atlas^23^ were averaged together within a subject and a paired t-test was used to test for differences between subtypes. All p-values were false discovery rate (FDR) corrected for multiple comparison.

### Behavioral Performance Differences Between Subtypes

A linear mixed-effect model was used to test for differences in accuracy, RT and confidence between subtypes. The model assessed the effects of subtype and condition with the subject as a random factor on behavioral performance.

### Brain-Behavior Relationship Within Each Subtype Compared to Across All Trials

To test if clustering trials improved the brain-behavior relationship estimation, we estimated the brain-behavior in each subtype and compared it to all trial pooled together. Specifically, we calculated the average activation in each brain region part of the Schaefer 200 brain atlas. Within each subject, for each region we correlated the trial activation strength with accuracy, RT and confidence for each subtype of trials and all trials together. We estimated the change in sensitivity in the brain-behavior relationship as:

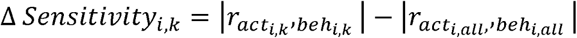

where |*r_acti,k_, _behi,k_* | is the absolute value of the correlation for trials in subtype *k* between activation strength in brain region i and behavioral performance and *|r_acti,all,_,_behi,all_* | is the absolute value in correlation between activation strength in brain region i and behavioral performance across all trials.

### SVM Classification

To corroborate our findings using modularity-maximization, we performed an analysis using Support Vector Machine (SVM) classifier using with MATLAB’s *fitcecoc.m.* For the analysis, we split the subjects in half and then repeated the clustering analysis on each half and trained an SVM classifier to predict the labels on the data from the other half of subjects. This analysis tests whether labels can be predicted on trials that were not included in the clustering. The SVM analysis was conducted with default parameters. Specifically, the SVM classifier utilized a linear kernel, with a 3^rd^ order polynomial function, a kernel offset of 0.1 for each element in the Gram matrix, and the prior distribution from each class is estimated from the relative frequencies of each class, Karush-Kuhn-Tucker complementarity conditions violation tolerance of 0.01.

### Transition Probabilities

We calculated transition probabilities by computing the probability of a trial from each subtype to be followed by a trial from any subtype.

### Data and Code Availability

The analysis was based on a combination of publicly available toolboxes, datasets and analysis specific scripts. Specifically, single-trial betas were estimated using GLMsingle and is available at https://github.com/cvnlab/GLMsingle. Clustering analysis was based on the Community Louvain part of the Brain Connectivity Toolbox (https://sites.google.com/site/bctnet/). Consensus clustering was determined using *consensus_iterative.m* (http://commdetect.weebly.com/). Unthresholded brain maps for each subtype are available at NeurVault, while behavioral data and analysis scripts are available at https://osf.io/kpnbs/.

## Acknowledgments

This work was supported by the National Institutes of Health (grant R01MH119189 to DR) and the Office of Naval Research (grant N00014-20-1-2622 to DR).

## Author contributions

Conceptualization: JN, DR

Methodology: JN, JY, NH, DR

Data Curation: JY, NH, JHK, SPK

Visualization: JN, DR

Funding acquisition: DR

Writing – original draft: JN, DR

Writing – review & editing: JN, JY, NH, JHK, SPK, DR

