## Supplemental Information for "Multiple Brain Activation Patterns for the Same Perceptual Decision-Making Task"

**Supplementary Results**

Control Analyses for the Subtypes

It is important to determine that the multiple activation patterns identified in each of the different tasks are not simply due to methodological choices, experimental factors, or noise in the data. As in Experiment 1, we found no clear relationship between subtypes and fMRI noise as measured with Frame Displacement, DVARS, or with each of the 6-motion parameters in Experiment 2 and Experiment 3 (**Fig. S4-S5**). To be as sensitive as possible, we conducted a series of pairwise comparisons between every two subtypes in every measure of subject motion. When using uncorrected tests, we obtained 2 significant differences from a total of 126 tests run, which is exactly equal to the 5% expected rate of significant results assuming no significant effects.

One methodological choice that might affect the results pertains to estimating the similarity between trials. The similarity between trails is estimated at the voxel-level, however, calculating similarity in high-dimensional voxel space can be unstable. Therefore, we performed the same analysis by first averaging all beta-values within an ROI part of the Schaefer 200 atlas^14^. Reflecting the voxel-wise analysis, the ROI analysis similarly identified 3 subtypes with consistent activation as those observed at the voxel Experiment 1, and Experiment 2 (**Fig. S6A, B)**. Further, using the ROIs to calculate the similarity increased sensitivity because for Experiment 3, the ROI analysis identified three subtypes which more closely reflected the other tasks compared to the two subtypes identified when the analysis was performed at the voxel-level (**Fig. S6C**). Critically, the ROI based analysis identified a subtype that exhibited increased activation in the DMN as was found in the voxelwise analysis.

Further, experimental factors at the trial level or demographics of the population may account for the different activation patterns. To determine if these confound account for the subtype, first, we assessed if the subtypes reflect changes over the course of the experiment as measured by the trial rank of each subtype. We found significant differences (P < 0.05 uncorrected) in the trial ranks among subtypes, but the difference subtypes were relatively small (**Fig. S7A**). Specifically, across all datasets, the largest difference in trial rank were 40 trials found in Experiment 1 (Subtype 1 = 355 trials; Subtype 2 = 338; Subtype 3 = 378). Note that if the subtypes occurred in the first third, middle and last third of the trials, then these numbers would be 117, 350, and 525. Instead in all subtypes, the trial rank values are closer to what is expected from a random assortment of 357 trials. The same analysis in the other datasets yielded even smaller differences in trial ranks: 8 trials for Experiment 2; and 5 trials for Experiment 3. As was found in Experiment 1, the differences were small and close to what would be expected by chance suggesting that the subtypes do not simply reflect changes that might be associated with slow process such as drift or learning over the course of the task.

Additionally, we assessed if the subtypes reflected differences in inter-trial interval (IIT). We found significant differences (P < 0.05 uncorrected) in IIT but the differences in Experiment 1-3 were again relatively small (**Fig. S7B**). Specifically, for Experiment 1, we found that the average IIT values for Subtypes 1, 2, and 3 were 1.00, 0.97, and 1.05 sec, respectively, which are close to the expected value of 1.00 sec. Similarly, differences in IIT were small and close to the expected value 2.00 sec in Experiment 2 (Subtype 1 = 1.95 sec; Subtype 2 = 2.01 sec; Subtype 3 = 2.06 sec). Additionally, in Experiment 3, the difference in the IIT were small and close to the expected value 5.50 sec values (Subtype 1 = 5.52 sec; Subtype 2 = 5.49 sec). These results indicate that methodological choices were not a significant factor in determining the composition of the subtypes.

Finally, we tested if there was an association between the subtypes and the demographics of the population (sex or age). We found no significant difference (P > 0.05) in the proportion of trials of each subtype between male and female participants for all four experiments (**Fig. S8A**). Further, we investigated if there were any associations between proportions of trials per subtype with age. For Experiments 1-3 we found no significant correlation between proportions of trials per subtype and age (P > 0.05; **Fig. S8B-D)**. These results indicate that age and sex were not significant factor determining the composition of subtypes. Taken together, these results suggest that the multiple activation patterns are not simply driven by trivial experimental factors, subject characteristics to various types of noise or experimental factors that might affect trial-by-trial activation patterns over the course of a task.

Transition Probabilities Between Subtypes

We estimated the transition probabilities between subtypes. If the same subtype is more likely to occur from trial to trial, then this could indicate that the subtypes may reflect infra-slow oscillations such as quasi-periodic patterns^1^ or changes in cognitive strategies^2^ that may lead to clusters of trials exhibiting similar activation patterns. In Experiment 1, 2 and the Working Memory task, the transition probabilities exhibited strong re-occurrence of a subtype suggesting slow infra-slow oscillations in brain activity or cognitive strategies may be associated with the different activation patterns (**Fig. S9**). However, in Experiment 3, the probability of a trial being part of the same subtype or transitioning to another subtype was almost equivalent suggesting that for this experiment infra-slow oscillations were not a factor. These results suggest that infra-slow oscillations such as quasi-periodic patterns or changes in cognitive strategies that may lead to clusters of trials exhibiting similar activation patterns.

Computational Modeling

We investigated a possible mechanism that would generate the multiple activation patterns from the structural connectivity and functional connectivity since both have been shown to be capable of predicting brain activation ^3,4^. The critical idea behind the model is that observed activations in each network arise from two factors: (1) a direct stimulus drive to the network, (2) indirect contributions from other brain networks that are the product of their own stimulus drive times the strength of their structural or resting-state functional connectivity with the given network (see supplemental methods for details; **Fig. S10A-C**). Specifically, the model assumes that observed activation, $A_{i}$, in network $i$ can be expressed as:

$$A_{i}=\sum_{j=1}^{7} s_{j} \times w_{ij}$$

where $s_{j}$ is the stimulus drive to network $j$, and $w_{ij}$ is the structural or resting-state functional connectivity strength between networks $j$ and $i$. Note that the final activation for each network also depends on the within-network connectivity strength, $w_{ii}$. We empirically estimated the structural and resting-state functional connectivity from diffusion weighted imaging and resting-state fMRI data of 50 subjects from the Human Connectome Project^5^.

Focusing on Experiment 1, the model allowed us to recover the strength of the stimulus drive to each network. The obtained stimulus drives mostly mirrored the observed stimulus activations, but with several important exceptions when using the structural connectivity (**Fig. S10D**). We found a very strong stimulus drive in VAN for Subtype 2, which suggests that one of the critical distinguishing factors of Subtype 2 may have been the presence of strong exogenous attention and saliency detection ^6–8^. Further, DAN showed consistently negative stimulus drive for all three subtypes even though it exhibited positive activations for Subtypes 1 and 2. These results indicate that the model can be used to track the complex interactions across networks.

It is worth noting that results are in part counter-intuitive because the DAN had a negative stimulus drive values for all three subtypes even though it exhibited positive activations. Nevertheless, this negative stimulus drive can be interpreted as stemming from the fact that the stimulus drive by itself does not contains any information regarding the task-relevant component or for producing the appropriate behavioral response, which are thought to be key elements of DAN’s cognitive function^9^. These elements are determined after the stimulus is processed by other networks such as the visual system and fed into the DAN. Thus, the input into the DAN may be strong enough to result in an increase in activation.

In addition, we compared the model to an alternative where communication between networks is dependent on the strength of the resting-state functional connectivity. However, when using the functional connectivity, all the stimulus drives values were negative even though various networks exhibited increased in activation (**Fig. S10E**). These negative stimulus drive values resulted from the resting-state functional connectivity containing both positive and negative in the matrix.

Therefore, we first made all resting-state functional connectivity values positive by taking the absolute value and re-ran the analysis. With only positive values, the stimulus drives reflected the observed activation patterns (**Fig. S10F**). Moreover, the model suggested a dissociation between brain networks with the strongest stimulus-drive and activation (**Fig. S11**). It is beyond the scope of this analysis to determine whether the resting-state functional connectivity should contain only positive values, but our results with the resting-state functional connectivity further highlight an existing issue pertaining to interpretation and estimation negative values in the resting-state functional connectivity values^10,11^.

Nonetheless, inferring a causal relationship from diffusion-based structural or functional connectivity has some limitations. Casual relationships are difficult to deduce from both structural and functional connectivity because the connections are undirected. Additionally, a recent study highlights that brain activity is depended on the geometry of the brain^12^. Taken together, these different findings highlight that brain activity can arises from complex interactions between structural connections, functional connections and/or geometry.

Multiple distinct subtypes of trials during working memory task

We also examine if multiple activation patterns exist in a different cognitive task. Specifically, we analyzed the *n*-back task data from the Human Connectome Project where subjects completed equal number of 0- and 2-back trials^5^. In the same manner as for Experiments 1-3, for each trial we estimated the task-induced brain activity in each voxel and pooled trials across subjects and *n*-back conditions, and then clustered the trials using modularity-maximization. Clustering identified three subtypes (**Fig. 12A**) present in both the 0- and 2-back conditions, but Subtype 1 was primarily in 0-back and Subtype 2 was primarily in the 2-back condition (**Fig. 12B**).

Examining the brain activation patterns, the visual network exhibited consistent activation in all three subtypes at the voxel (**Fig. 12C**) and among large-scale brain network (**Fig. 12D**). Further, as in Experiments 1-3, the subtypes were consistent across a range of gamma values (0.8 to 1.1; **Fig. 12E**) and the SVM classifier correctly identified 75.5% of trials (**Fig. 12F**). Further, we examined if there were behavioral differences between subtypes using a mixed-effect model as in the working memory task and we found significant differences in accuracy (t(3837) = 2.74, P = 0.006; **Fig. S12H**) but not for RT (t(3837) = 0.23, P = 0.82; **Fig. S12H**).

Additionally, we performed the analysis within each condition separately to ensure that the subtypes do not represent differences between the two *n*-back conditions, and we found the same patterns of activation within the 0- and 2-back separately (**Fig. S13**). Moreover, as in Experiment 1-3, the same subtypes were observed when the analysis was conducted at the ROI level (**Fig. S14A, B**), minor differences in trial rank (**Fig. S14C**) and no relationship sex **Fig. S14D**). However, motion associated noise was a factor in the n-back task (**Fig. S15**).

Overall, the activation pattern observed between subtypes did not exhibit strong differential activation that was observed in Experiment 1, 2 and 3. This finding could reflect the higher level of fMRI noise, particularly z-motion which could make it difficult to identify the subtypes. Furthermore, the n-back task contains 80 trials per subject compared to the hundreds of trials in the Experiment 1, 2 and 3 which adds to the challenge of identifying subtypes. Nonetheless, there may be tasks that do not contain multiple patterns of activation, but we do not want to make strong claims about subtypes not existing all n-back datasets, especially because of the limitations of the HCP data (larger motion and fewer trials).

**Supplementary Methods**

Model Specification

We sought to characterize the hidden stimulus drive to each of the seven brain networks by fitting a computational model to the data. The model generates brain activity among large-scale brain networks using minimal assumptions. This approach is inspired by prior work on examining the influence interactions between large-scale brain networks have on the activity of an individual brain network^13,14^. Structural connectivity has been shown that it can be used to predict brain activity and thus allowing us to characterize how the observed activation patterns might arise from a stimulus drive and network interactions^4,15^. Beside structural connectivity, task dependent brain activity can be simulated from the resting-state functional connectivity^3^.

The critical idea behind the model is that observed activations in each network arise from two factors: (1) a direct stimulus drive to the network, (2) indirect contributions from other brain networks that are the product of their own stimulus drive times the strength of their structural connectivity with the given network. Specifically, the model assumes that observed activation, $A_{i}$, in network $i$ can be expressed as:

$$A_{i}=\sum_{j=1}^{7} s_{j}\times w_{ij}$$

where $s_{j}$ is the stimulus drive to network $j$, and $w_{ij}$ is the structural connectivity strength between networks $i$ and $j$. Note that the final activation for each network also depends on the within-network connectivity strength, $w_{ii}$. We can obtain the stimulus drive (“s”) by *s* = *A* $\times$ *inv*(*w*), where *inv* is the inverse of the matrix. The model focuses on the group-level activation since each subtype was present in all subjects. Specifically, A_i_ represents the observed activation in the seven large-scale networks part of the Schaefer atlas^14^. A_i_ is estimated by averaging estimated activation values from all voxels associated with each network all trials as a result there are no time dependencies in the model. For visualization purposes $s_{j}$ is divided by 1000.

Structural Connectivity

Structural connectivity between brain regions was estimated from 50 healthy human data from the S1200 release from the Human Connectome Project (HCP)^16^. Fiber tracking was conducted using DSI Studio with a modified FACT algorithm^17^. Diffusion MRI images were reconstructed in MNI space using generalized q-sampling imaging and fiber tracking was performed until 250,000 streamlines were reconstructed with angular threshold of 50^o^_,_ step size of 2 mm, minimum length of 10mm, and maximum length of 400mm.

For each subject, an undirected weighted structural connectivity matrix, $w$, containing 200 brain regions part of the Schaefer atlas was constructed by counting the number of streamlines between brain regions. Connectivity matrices were normalized by dividing the number of streamlines (*T*) by the combined volumes (*v*) of regions *i* and *j*:

$$W_{ij}= \frac{T_{ij}}{v_{i}+v_{j}}$$

The strength of the connectivity between large-scale brain networks was estimated by first averaging the matrices, $w$, across subjects to generate a group structural connectivity map followed by averaging the strength of the connectivity between regions between the seven large-scale brain networks in the Schaefer Atlas. This resulted in a 7x7 matrix where each entry represents the strength of the connectivity within and between brain networks.

Resting-State Functional Connectivity

Resting-state functional connectivity between brain regions was estimated from 48 healthy human data from the S1200 release from the Human Connectome Project. For each subject, the resting-state functional connectivity was estimated using the Pearson correlation between the timeseries date among the 200 brain regions part of the Schaefer 200 atlas. A group 7x7 matrix was then estimated my first averaging the individual subject-level resting-state functional connectivity and further averaging the correlation values among the seven large-scale networks associated with Schaefer atlas. However, estimating the resting-state functional connectivity in this manner produces negative values and there is some disagreement on how to interpret these negative values^10,11^. Therefore, we additionally, used an estimate of the resting-state functional connectivity with all positive values in the model by first taking the absolute value.

HCP Working Memory Task

The analysis utilized the minimally preprocessed n-back data collected part of the Human Connectome Project^5^. The HCP n-back analysis was based on 48 subjects in order to equate the number of subjects in Experiment 1. (Note, we downloaded 50 subjects but for two subjects the data was missing.) Specifically, the analysis utilized WM_RL data in which each subjects completed 40 trials of 0- and 2-back trials. MRI scanning was done using a customized 3 T Siemens Connectome Skyra using a standard 32-channel Siemens receive head coil and a body transmission coil. T1-weighted high resolution structural images acquired using a 3D MPRAGE sequence with 0.7 mm isotropic resolution (FOV = 224 mm, matrix = 320, 256 sagittal slices, TR = 2400 ms, TE = 2.14 ms, TI = 1000 ms, FA = 8°). Working Memory fMRI data were collected using gradient-echo echo-planar imaging (EPI) with 2.0 mm isotropic resolution (FOV = 208 × 180 mm, matrix = 104 × 90, 72 slices, TR = 720 ms, TE = 33.1 ms, FA = 52°, multi-band factor = 8, 1200 frames, ~15 min/run).

Prior to estimating the task-induced brain activity on individual trials, the fMRI files were spatial smoothed with 10 mm full width half maximum (FWHM) Gaussian kernel. Task-induced brain activity in each voxel was estimated using GLMsingle^18^. Additionally, global signal and motion parameters were included as factors in GLMsingle. For clustering, we pooled trials across subjects and *n*-back conditions (N_subj_ = 48; N_trials_ = 3840) in the same manner as in the perceptual decision-making tasks.

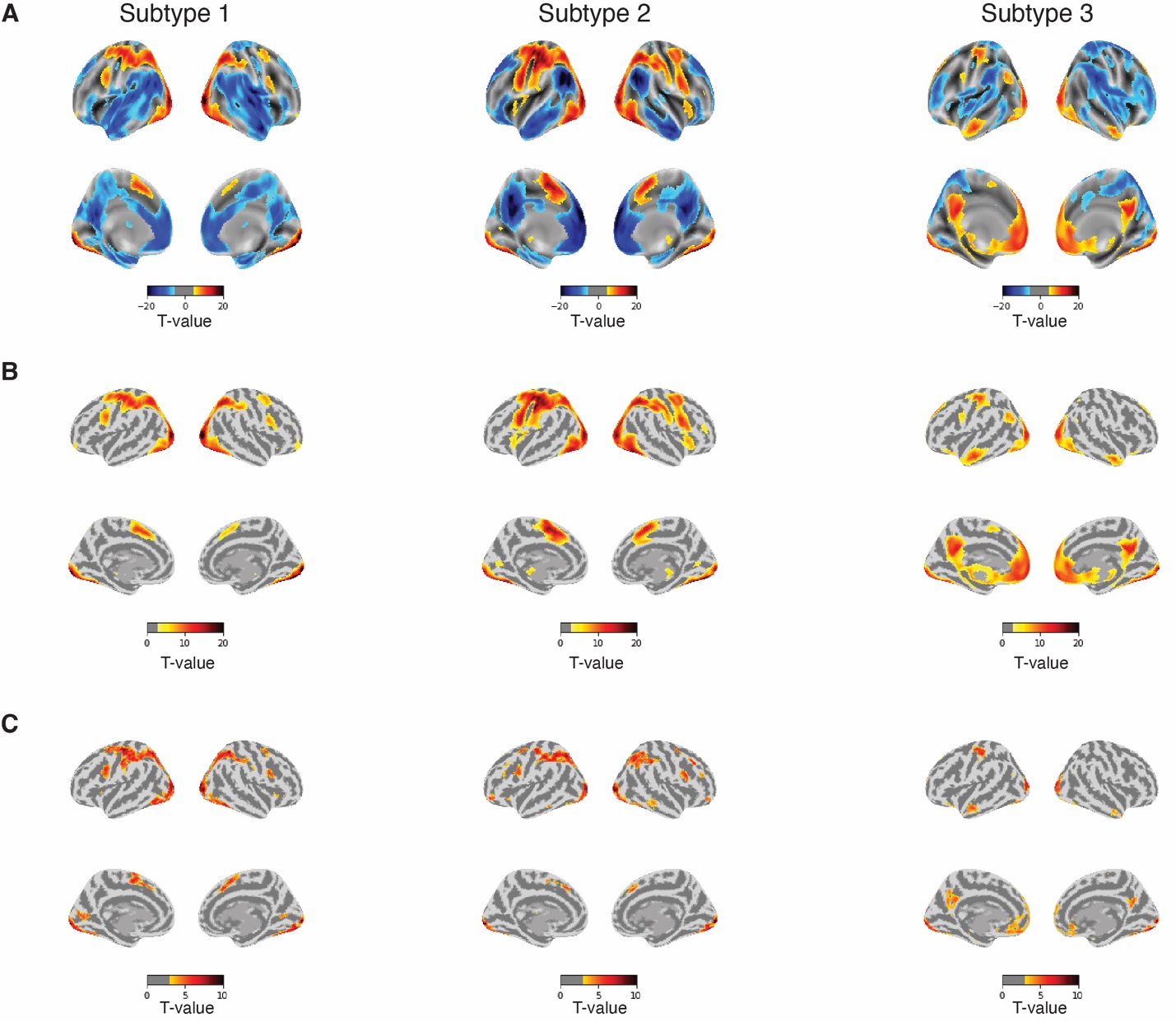

**Figure S1. Standard GLM analysis with trial subtype labels as factors in the regression in Experiment 1.** Traditional GLM analysis was able to reproduce the changes in brain activity for (A) Subtype 1, (B) Subtype 2, and (C) Subtype 3. Brain maps are thresholded at FDR-corrected p < 0.01. (B) Cluster-corrected analysis results of brain maps from panel A. To identify brain regions that increased in activation, brain maps were first thresholded at p < 0.001 uncorrected followed by cluster-based correction. (C) Clustering of fMRI data smoothed with 4mm full width half-max Gaussian kernel identified three subtypes. Brain maps were first thresholded at p < 0.001 uncorrected and followed by cluster-based correction at p < 0.05.

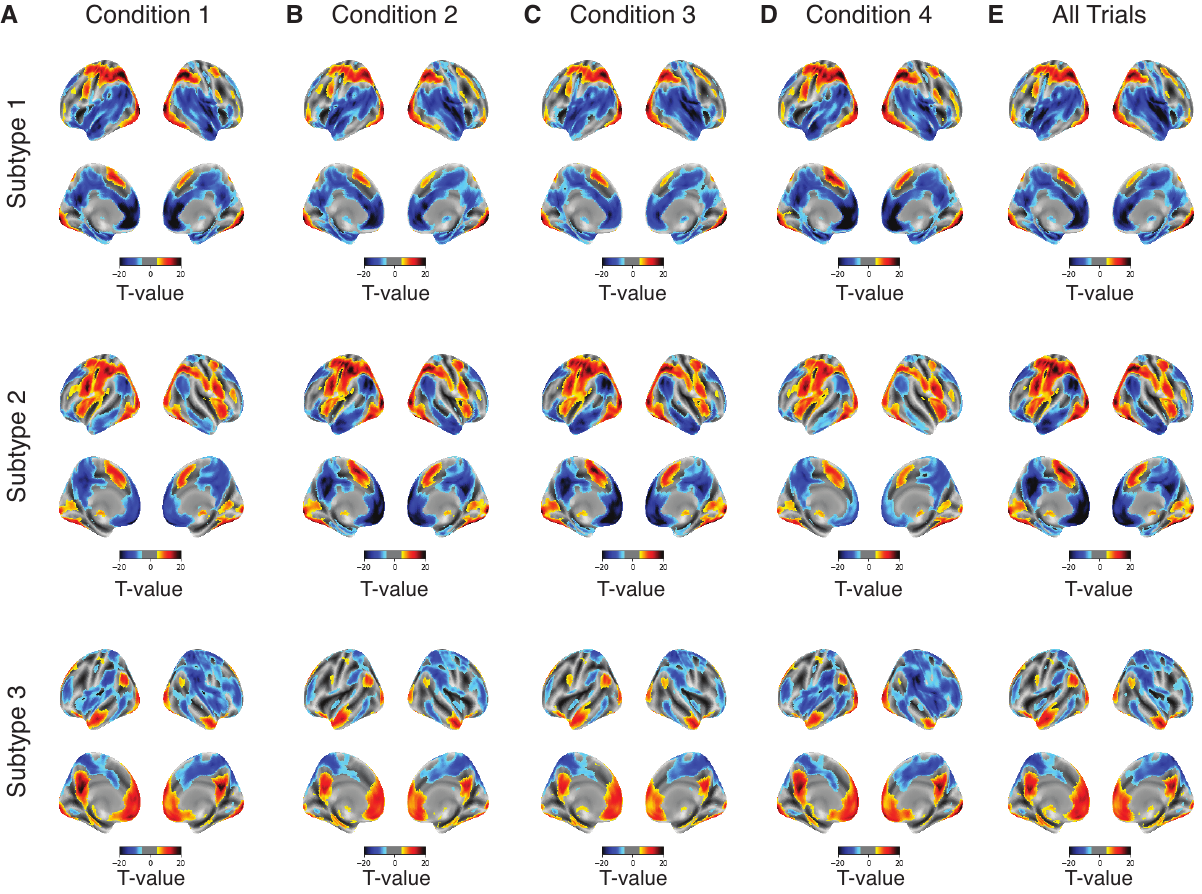

**Figure S2. Subtypes are present within each condition in Experiment 1.** Modularity-maximization based clustering identified three subtypes of trials within each condition in Experiment 1. Activation patterns in (A) Condition 1, (B) Condition 2, (C) Condition and (D) Condition 4, and (E) all trials pooled together. Each condition represents one of four different dot ratios were used – 80/60, 80/70, 100/80, and 100/90, where the two numbers indicate the number of dots from each color. Single-trial beta responses estimated with a general linear model (GLM) using GLMsingle^20^. We estimated the similarity across the activations between pairs of trials using Pearson correlation and clustered all trials using modularity-maximization to identify consistent activation patterns^21^. Panel E is the same as in Figure 2 and is included for comparison. Brain maps are thresholded at P_FDR-corrected_ < 0.01.

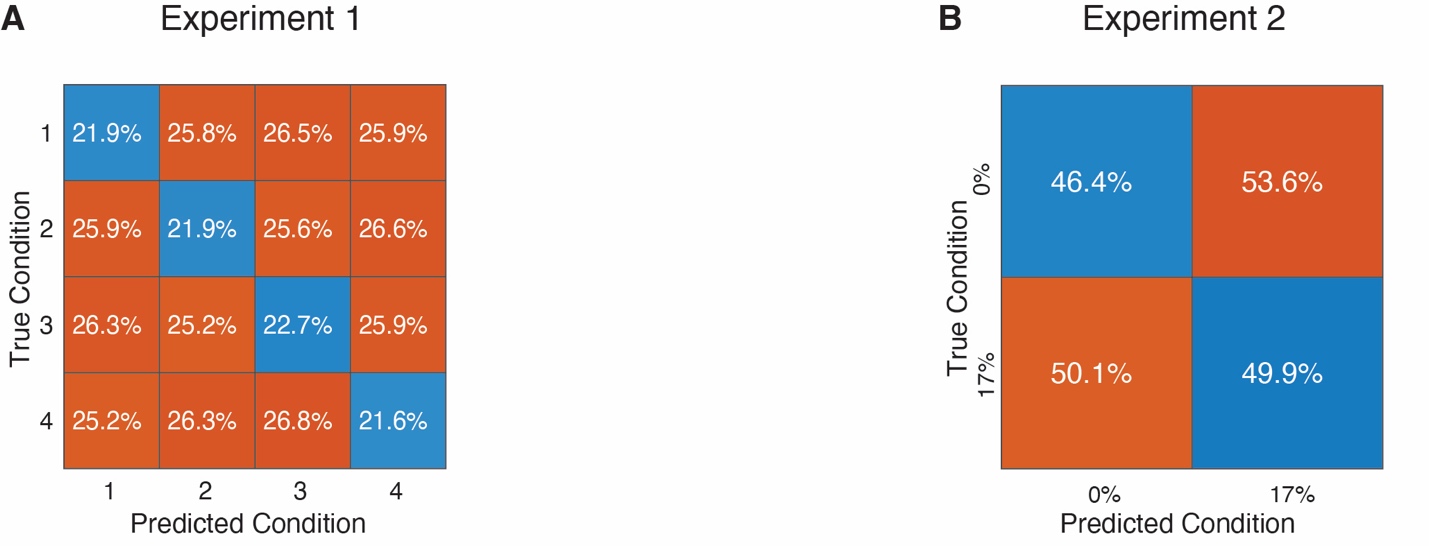

**Figure S3. Accuracy of SVM classification of task conditions of Experiment 1 and 2.** (A) SVM classifier to predict the condition labels in Experiment 1. An SVM model was trained on data from half the of the subject and tested on the data from the remaining subjects. (B) Same as panel A, but for Experiment 2. In both Experiment 1 and 2, classification was below of close to what is expected at random – 25% and 50%, respectively. (Note for Experiment 3, the task design did not contain conditions).

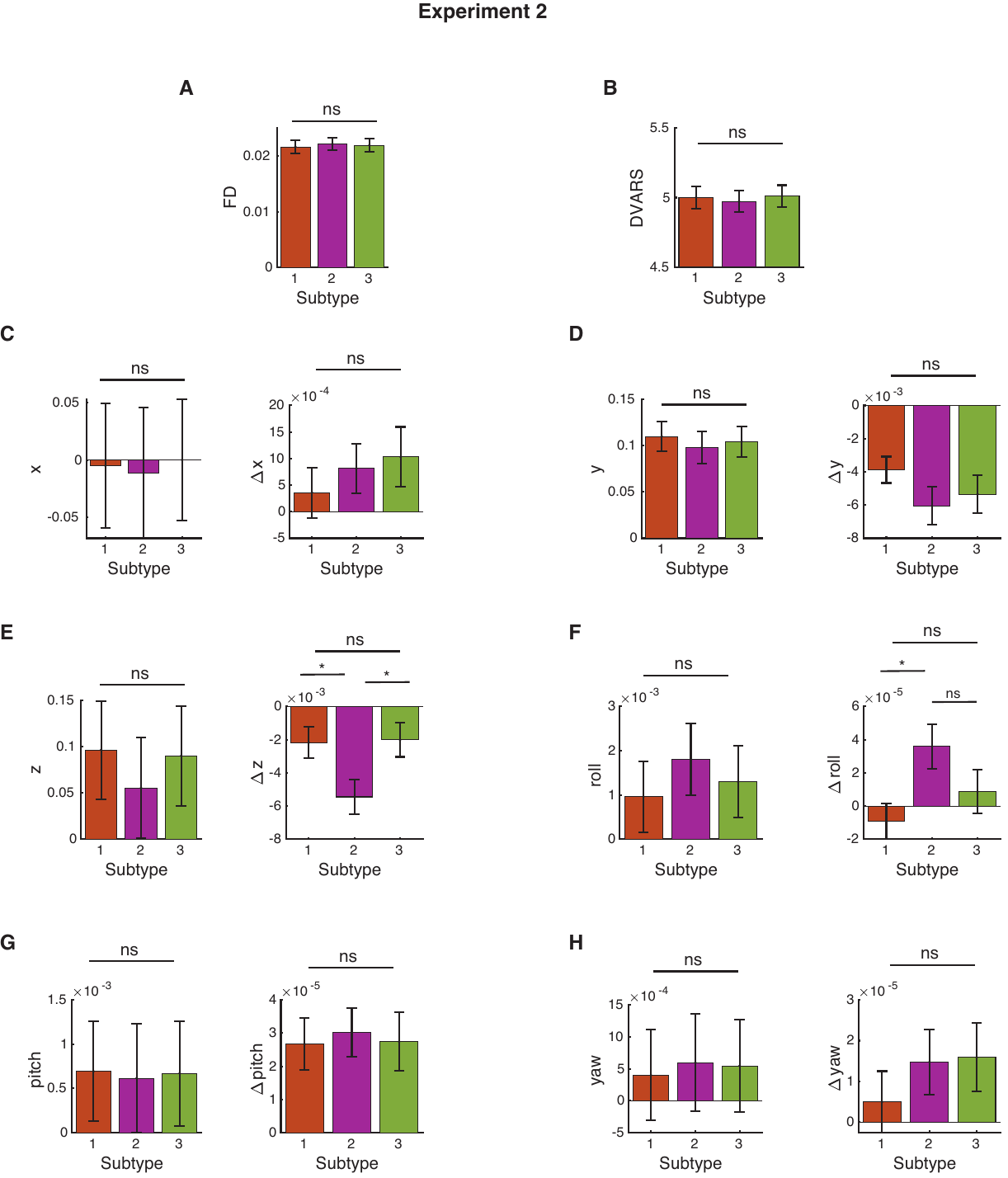

**Figure S4. Differences in motion parameters for Experiment 2**. (A) Frame displacement (FD), (B) DVARS, (C) x-, (D) y-, (E) z-, (F) roll-, (G) pitch-, (H) yaw-direction. For each trial we estimated 14 different motion associated artifacts. Estimated motion values were averaged per subtype within a subject and statistical differences were determined using paired-samples t-test. For panels C-H, right panels show the 1^st^ derivatives. *, P < 0.05; ns, not significant

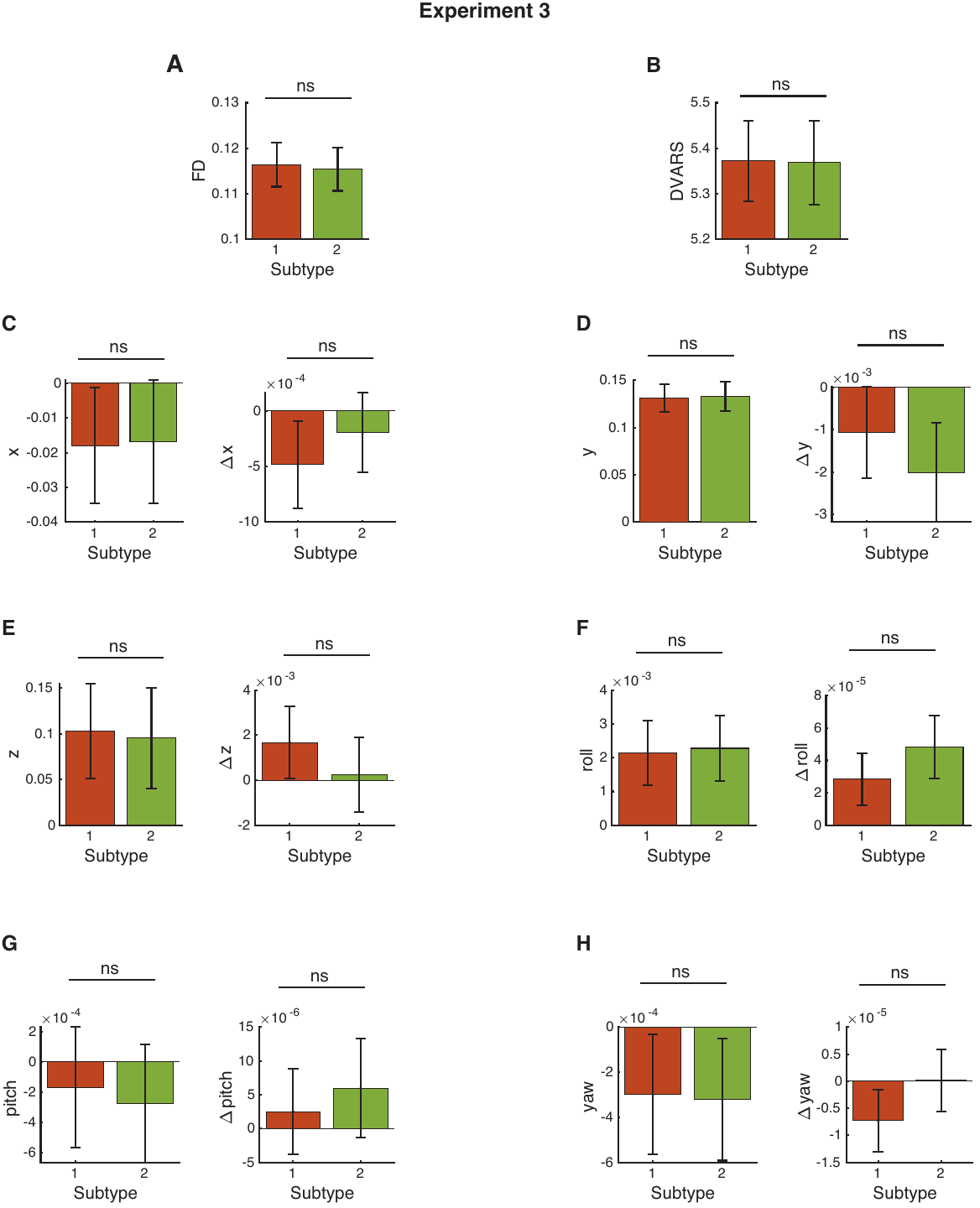

**Figure S5. Differences in motion parameters for Experiment 3**. (A) Frame displacement (FD), (B) DVARS, (C) x-, (D) y-, (E) z-, (F) roll-, (G) pitch-, (H) yaw-direction. For each trial we estimated 14 different motion associated artifacts. Estimated motion values were averaged per subtype within a subject and statistical differences were determined using paired-samples t-test. For panels C-H, right panels show the 1^st^ derivatives. ns, not significant.

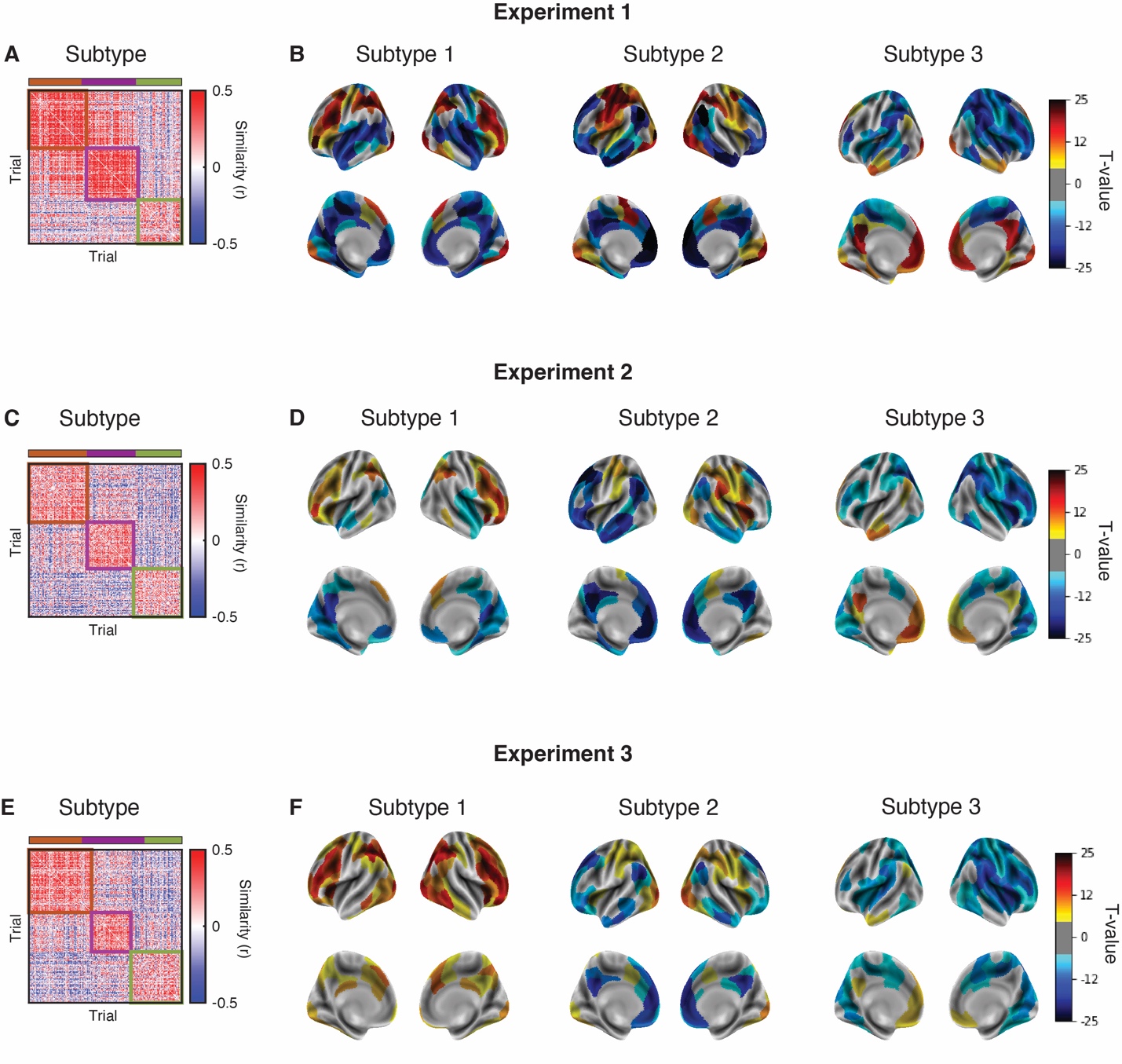

**Figure S6. ROI based clustering analysis.** Modularity-maximization based clustering on the activation from 200 brain regions part of the Schaefer atlas identified three subtypes of trials in (A, B) Experiment 1, (C, D) Experiment 2, and (E, F) Experiment 3. ROI-level activations were estimated by averaging all beta values from all voxels within a region. Pearson correlation was used to estimate the similarity between pairs of trials and clustered using modularity-maximization to identify consistent activation patterns^21^. Brain maps are thresholded at P_FDR-corrected_ < 0.01.

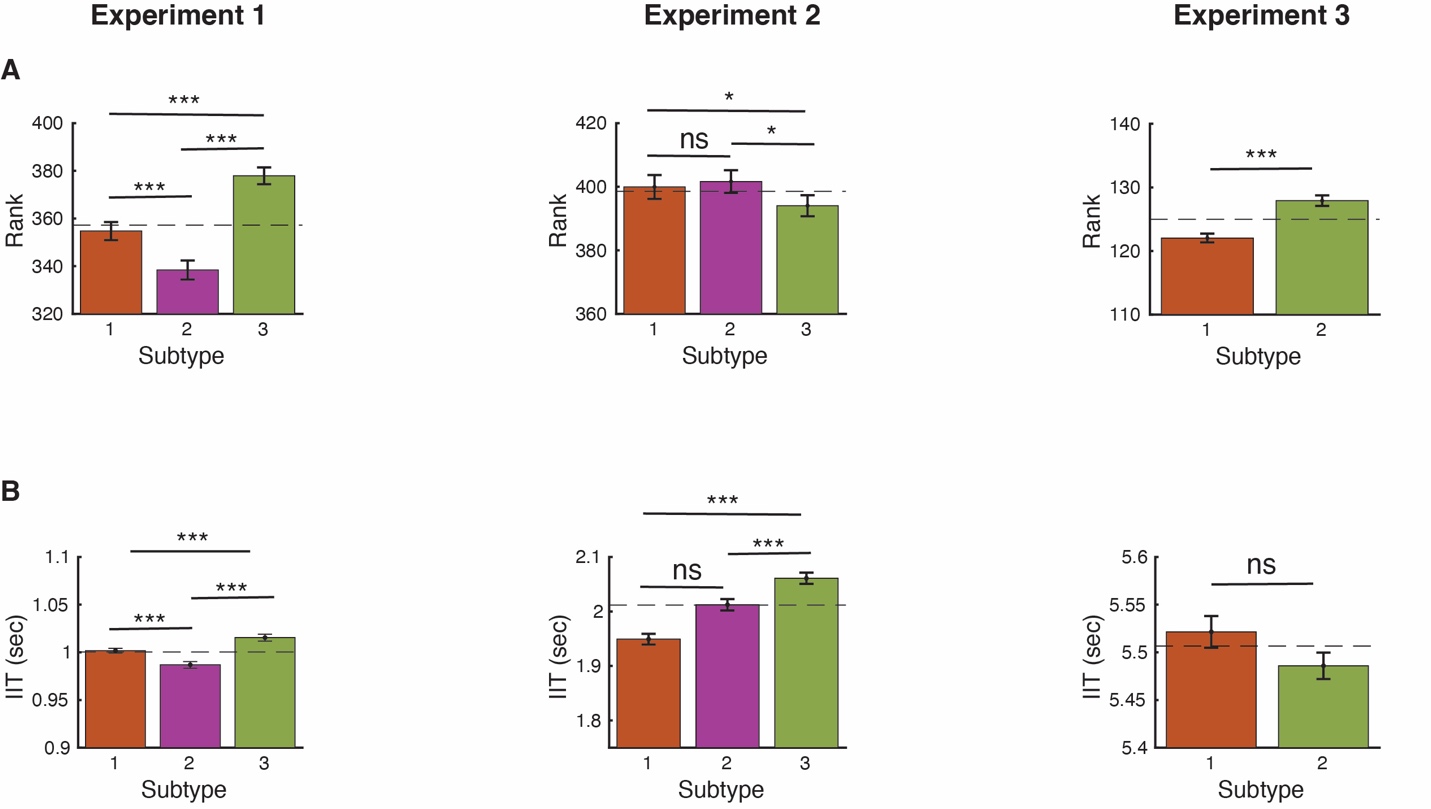

**Figure S7**. **Subtype trial rank and inter-trial interval across subtypes and tasks**. (A) Differences in trial rank per subtype to determine if the subtypes reflect slow changes over the course of the experiment. (B) Difference in inter-trial interval (IIT). IIT is the time after the response in the current trial to the onset of the stimulus in the next trial. Assessing the difference in IIT determines if the subtypes reflect changes in brain activity occurring on trials with longer intervals. The dashed line corresponds to the expected value from a random. Note that for the working memory task the IIT was fixed between trials. *** p < 0.001; * p < 0.05; ns = not significant.

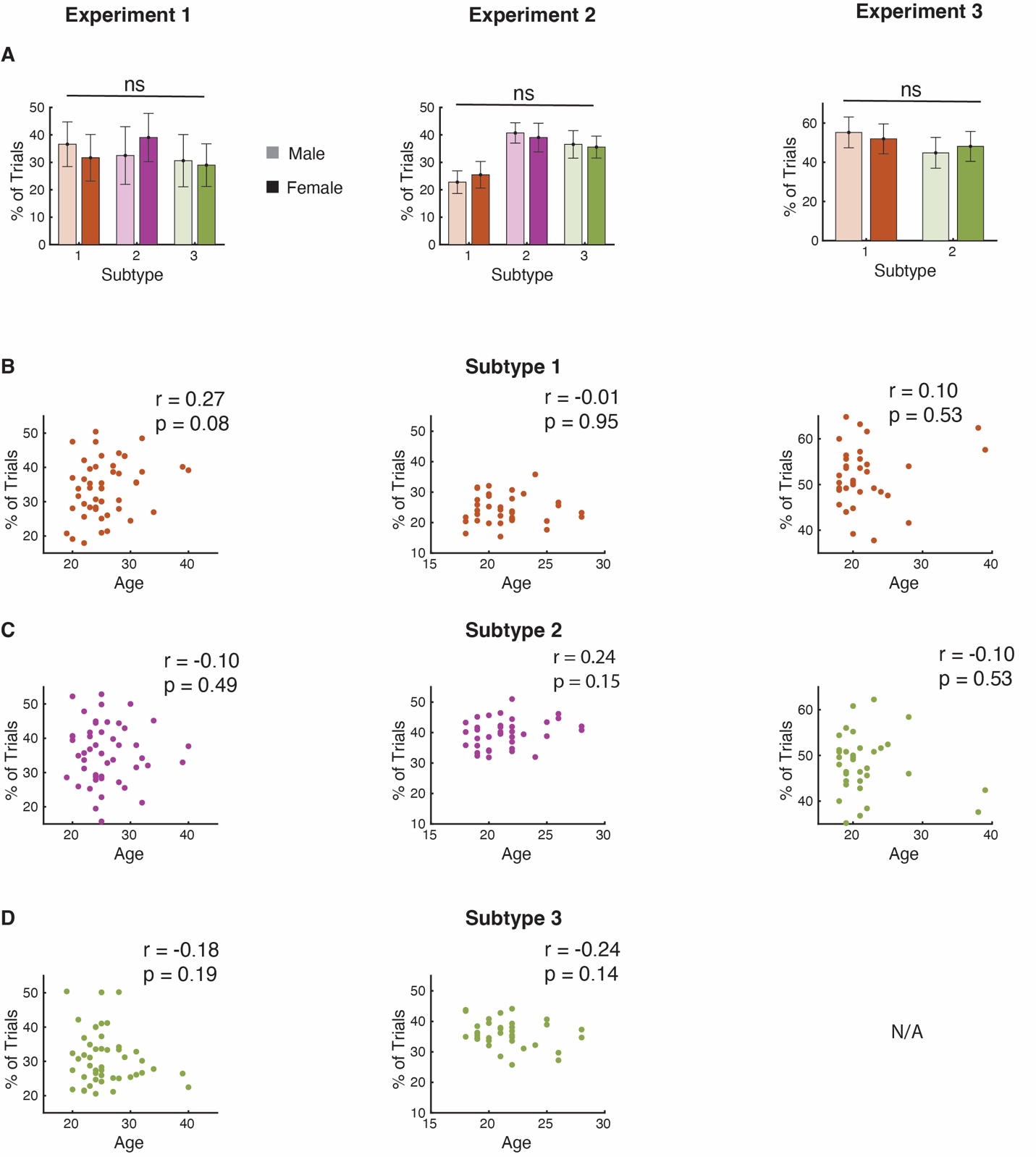

**Figure S8**. **Association between subtype with age and sex**. (A) No significant association between sex and the percentage of trials per subtype. For each subtype, differences in the proportion of trials were assessed between males and female subjects. Statistical differences were determined using independent samples t-test. (B-D) Correlation between age and percentage trials per subtype. Note that for Experiment 3, the analysis identified only two subtypes. ns, not significant.

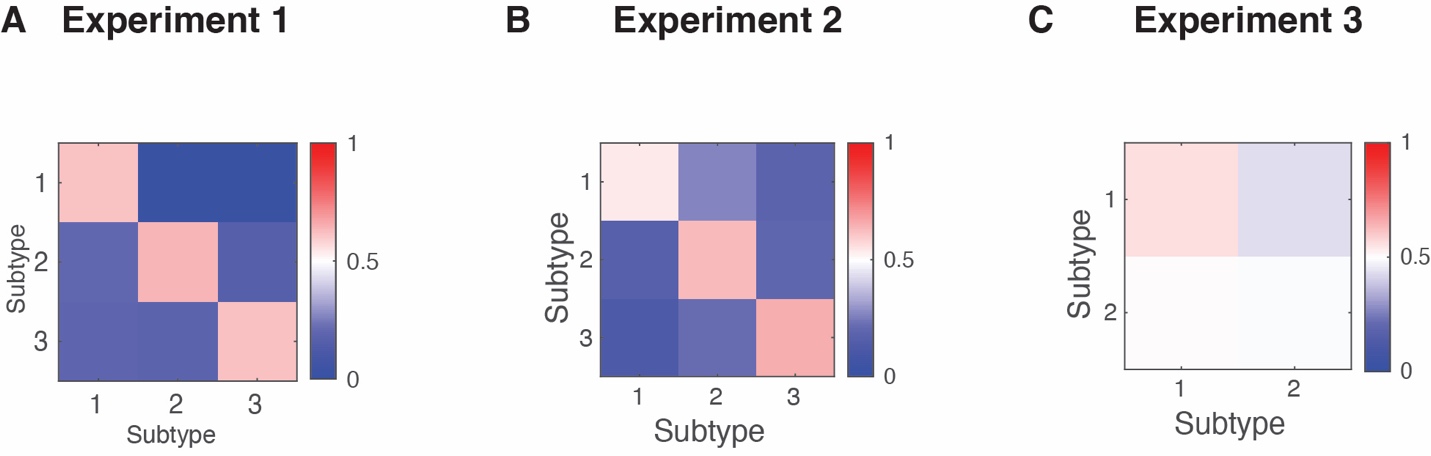

**Figure S9. State-transition matrix.** Transition probabilities among subtypes for (A) Experiment 1, (B) Experiment 2, and (C) Experiment 3. The transition probability represents the proportion of trial in which the subtype associated with each trial stayed the same between two consecutive trials or changed to one of the other subtypes. In Experiments 1/2, the transition probabilities strongly deviated from random (0.33). Whereas in Experiment 3, the transition probabilities were close to random (0.5).

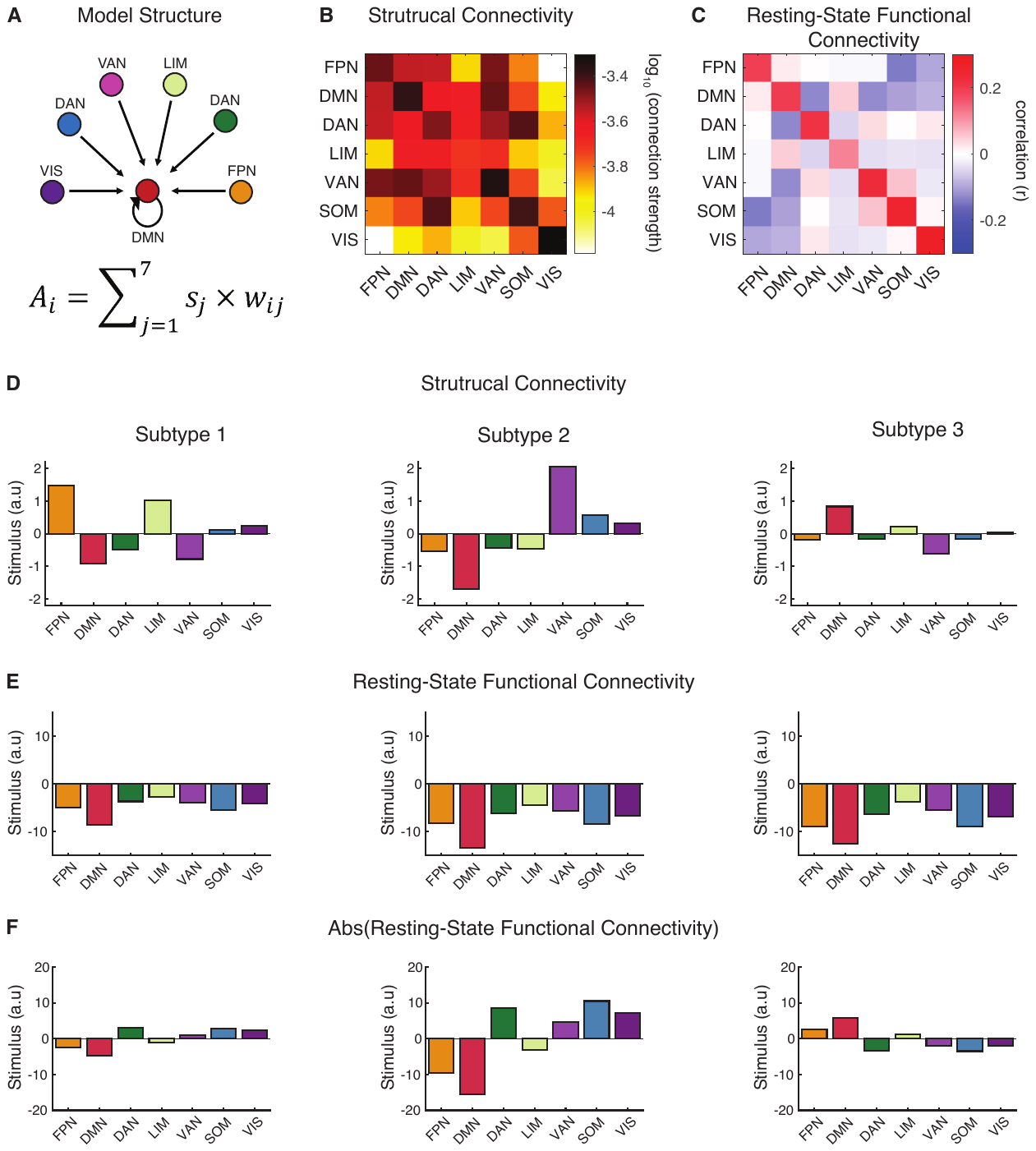

**Figure S10. Model structure and stimulus drive in Experiment 1.** (A) Graphical depiction of the model. Activation (*A_i_*) for a representative network, the DMN, is modeled as the product of the stimulus drive (*s_j_*) and the strength of the connectivity between the DMN and all other networks (*w_ij_*). (B) Group-level structural connectivity between 7 large-scale networks. (C) Resting-state functional connectivity from the same 7 large scale networks. The stimulus drive for each of the subtypes in Experiment 1 estimated from *w* set to the (D) structural connectivity, (E) RSN, or (F) absolute value of the RSN to remove the negative values. FPN, Frontal Parietal Network; DMN, Default Mode Network; DAN, Dorsal Attention Network; LIM, Limbic Network; VAN, Ventral Attention Network; SOM, Somatomotor Network; VIS, Visual Network.

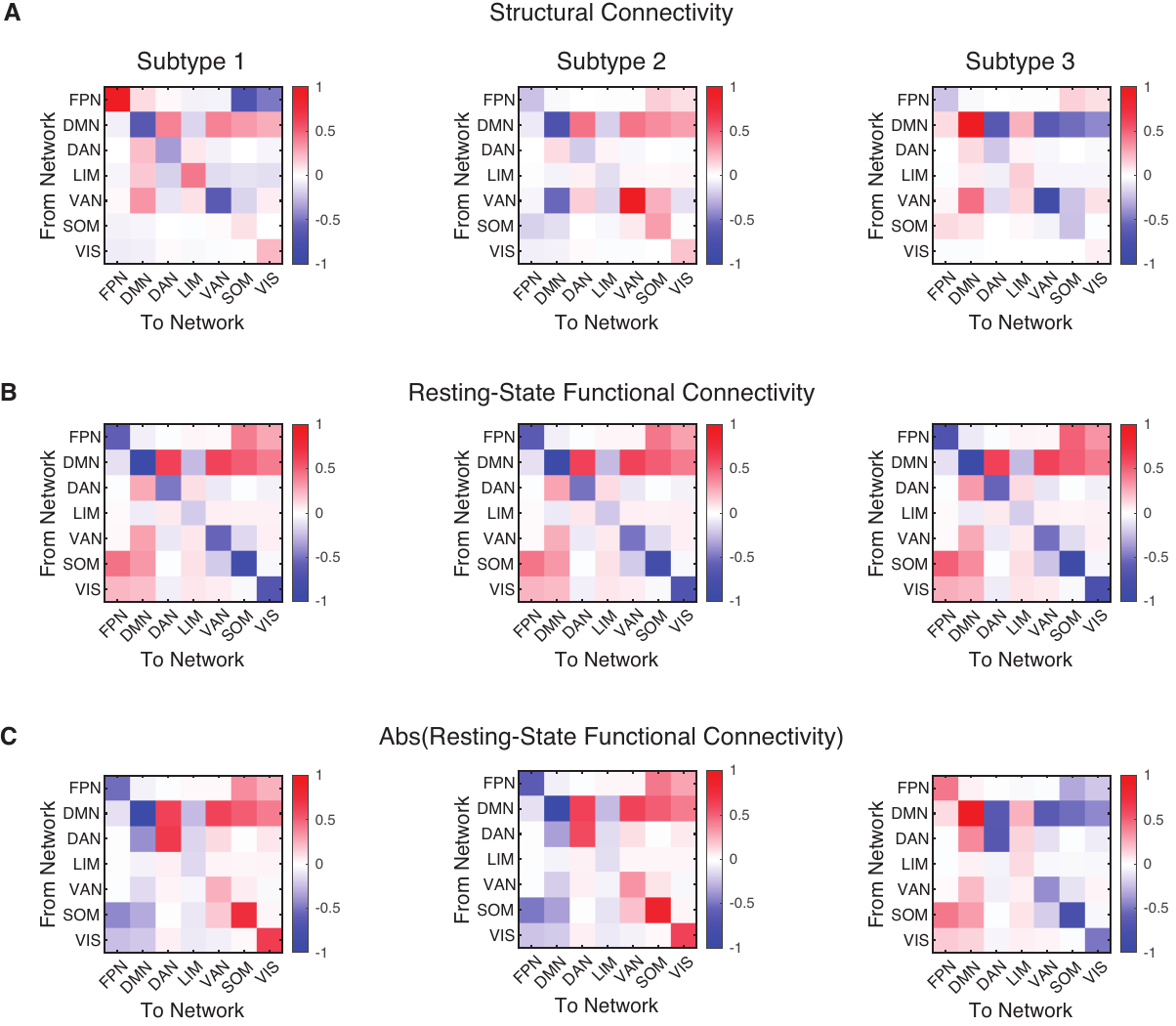

**Figure S11. Estimated interactions among large-scale brain networks in Experiment 1.** The effect of the stimulus drive modulated by connectivity strength between networks estimated when using (A) the structural connectivity, (B) resting-state functional connectivity, and (C) absolute value of the resting-state functional connectivity to remove the negative values. FPN, Frontal Parietal Network; DMN, Default Mode Network; DAN, Dorsal Attention Network; LIM, Limbic Network; VAN, Ventral Attention Network; SOM, Somatomotor Network; VIS, Visual Network.

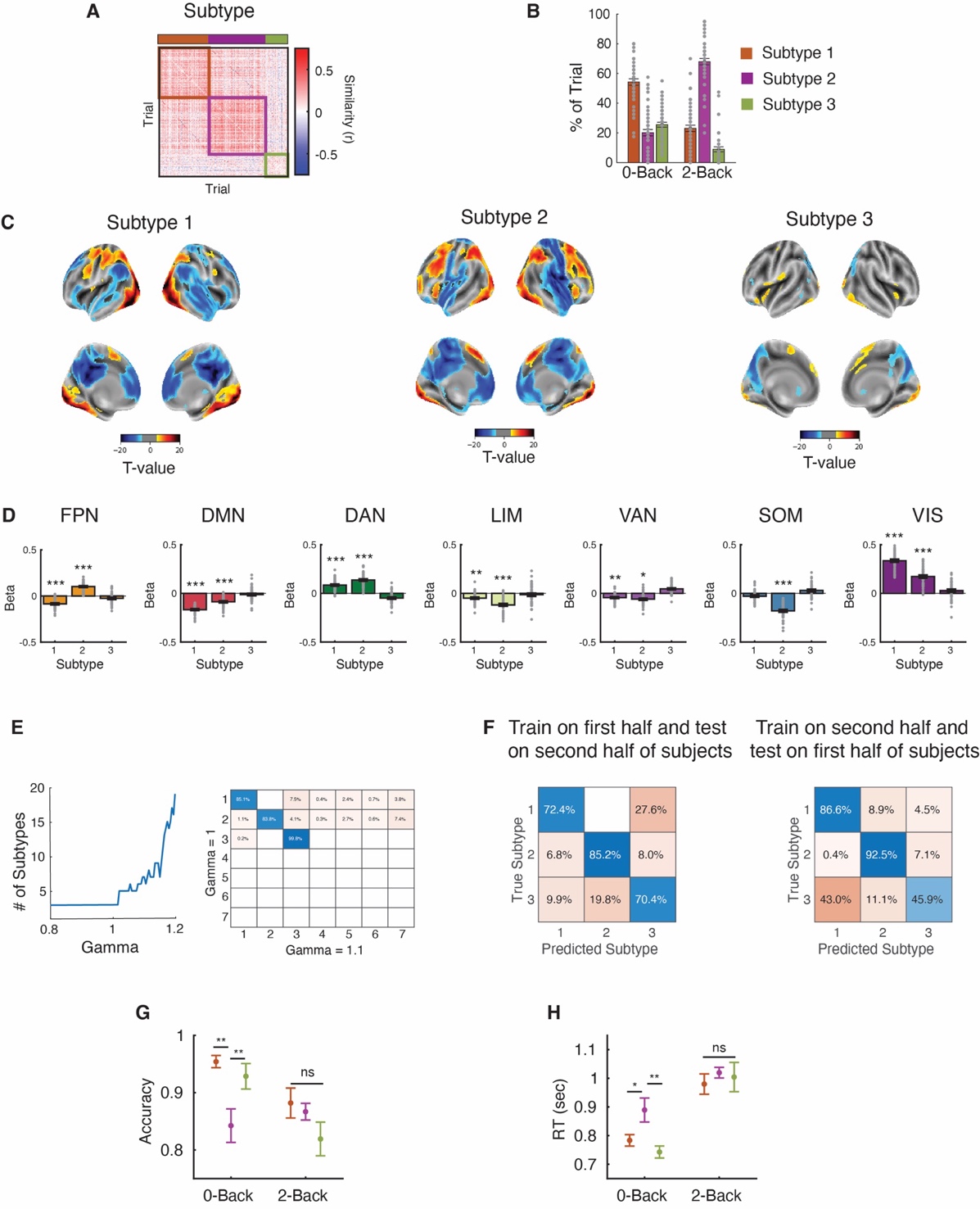

**Figure S12. Subtypes of trials in an *n*-back task from the Human Connectome Project dataset.** (A) Modularity-maximization clustering identified three subtypes of trials. (B) The percent of trials classified as Subtype 1, 2, or 3 in the 0- and 2-back conditions. Activation at the (C) voxel-level and (D) across large maps for each subtype. (E) Sensitivity of clustering to resolution parameter. Subtype were stable over a range of gamma, from 0.8 to 1.01 (*left*). We compared the clusters obtained with gamma = 1 and gamma = 1.1 (*right*). (F) SVM Classification. The SVM classifier correctly labeled on average 75.5% of trials across all tasks. (G, H) Differences in accuracy and reaction times between subtypes. *** P_FDR-corrected_ < 0.001; ** P_FDR-corrected_ < 0.01; * P_FDR-corrected_ < 0.05. FPN, Frontal Parietal Network; DMN, Default Mode Network; DAN, Dorsal Attention Network; LIM, Limbic Network; VAN, Ventral Attention Network; SOM, Somatomotor Network; VIS, Visual Network.

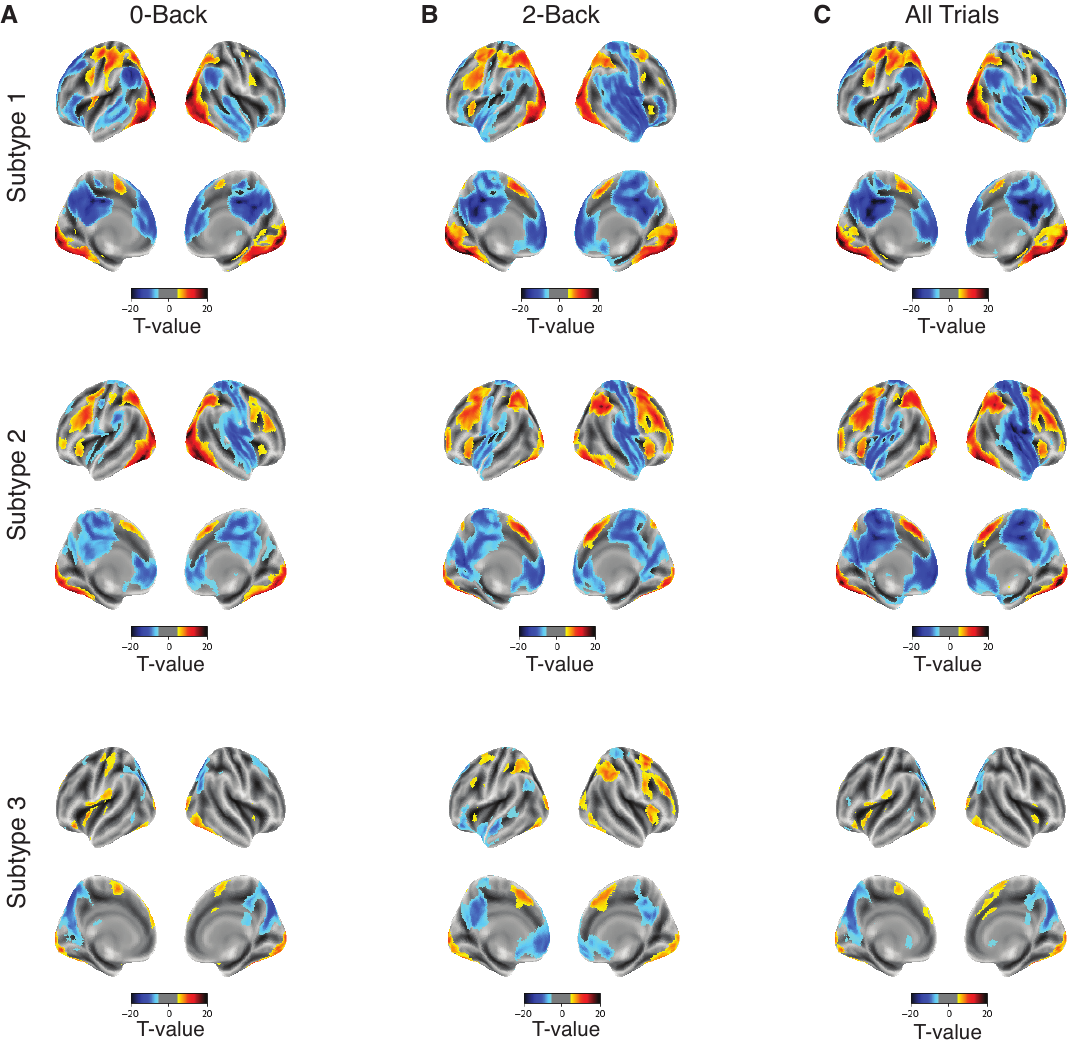

**Figure S13. Subtypes are present within each condition in the working memory task.** Modularity-maximization based clustering identified three subtypes of trials within(A) 0-Back, (B) 2-Back, compared to when pooling (C) all trials. Single-trial beta responses estimated with a general linear model (GLM) using GLMsingle^20^. We estimated the similarity across the activations between pairs of trials using Pearson correlation and clustered all trials using modularity-maximization to identify consistent activation patterns^21^. Panel C is the same as in Figure S6 and is included for comparison. Brain maps are thresholded at P_FDR-corrected_ < 0.01.

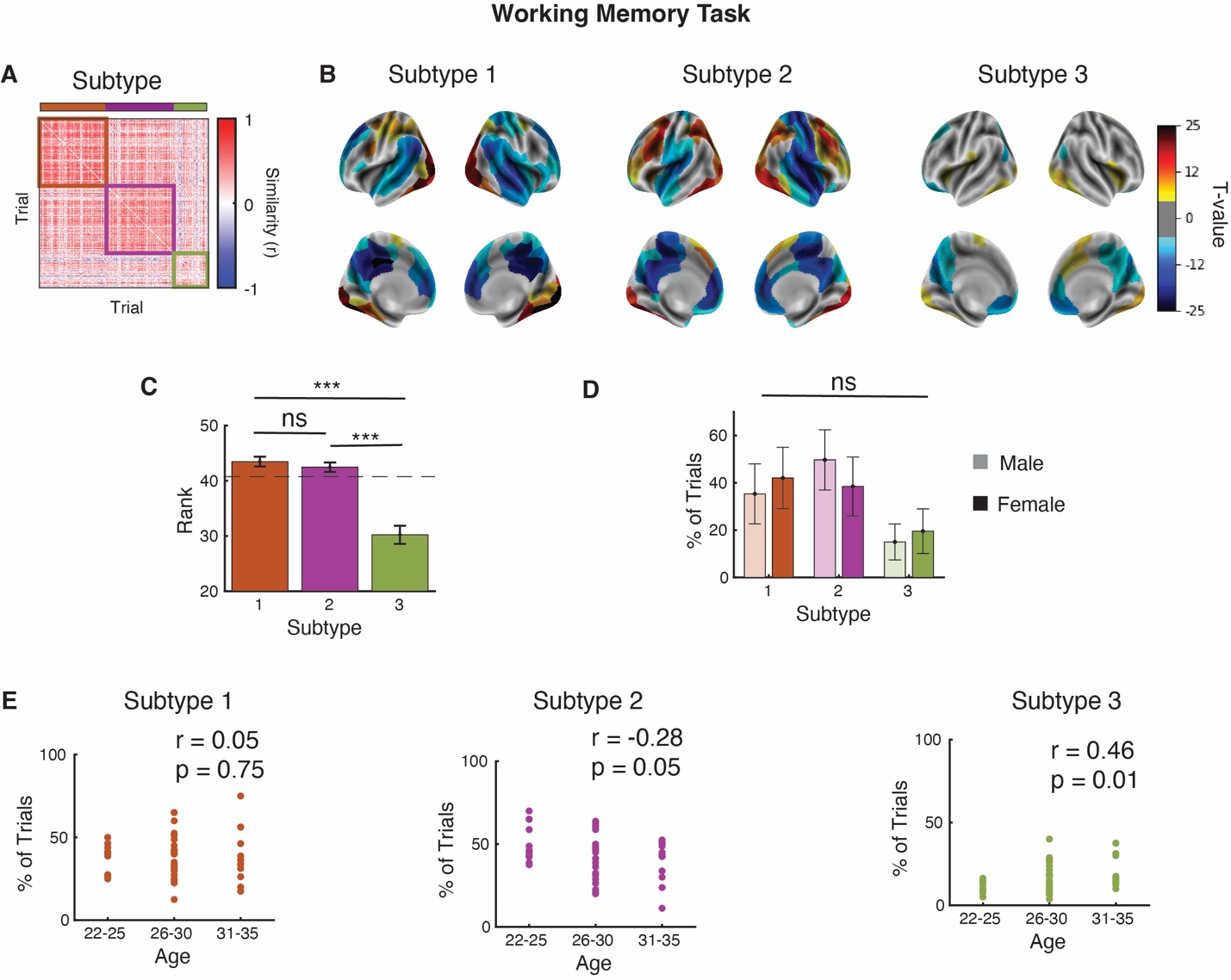

**Figure S14. HCP working memory.** (A, B) Modularity-maximization based clustering on the activation from 200 brain regions part of the Schaefer atlas identified three subtypes of trials.

(C) Differences in trial rank per subtype to determine if the subtypes reflect slow changes over the course of the experiment. (D) No significant association between sex and the percentage of trials per subtype. For each subtype, differences in the proportion of trials were assessed between males and female subjects. Statistical differences were determined using independent samples t-test. (E) Correlation between age and percentage trials per subtype. Note that caution is warranted given the relatively small number of subjects (N_subj_ = 40) in the sample.

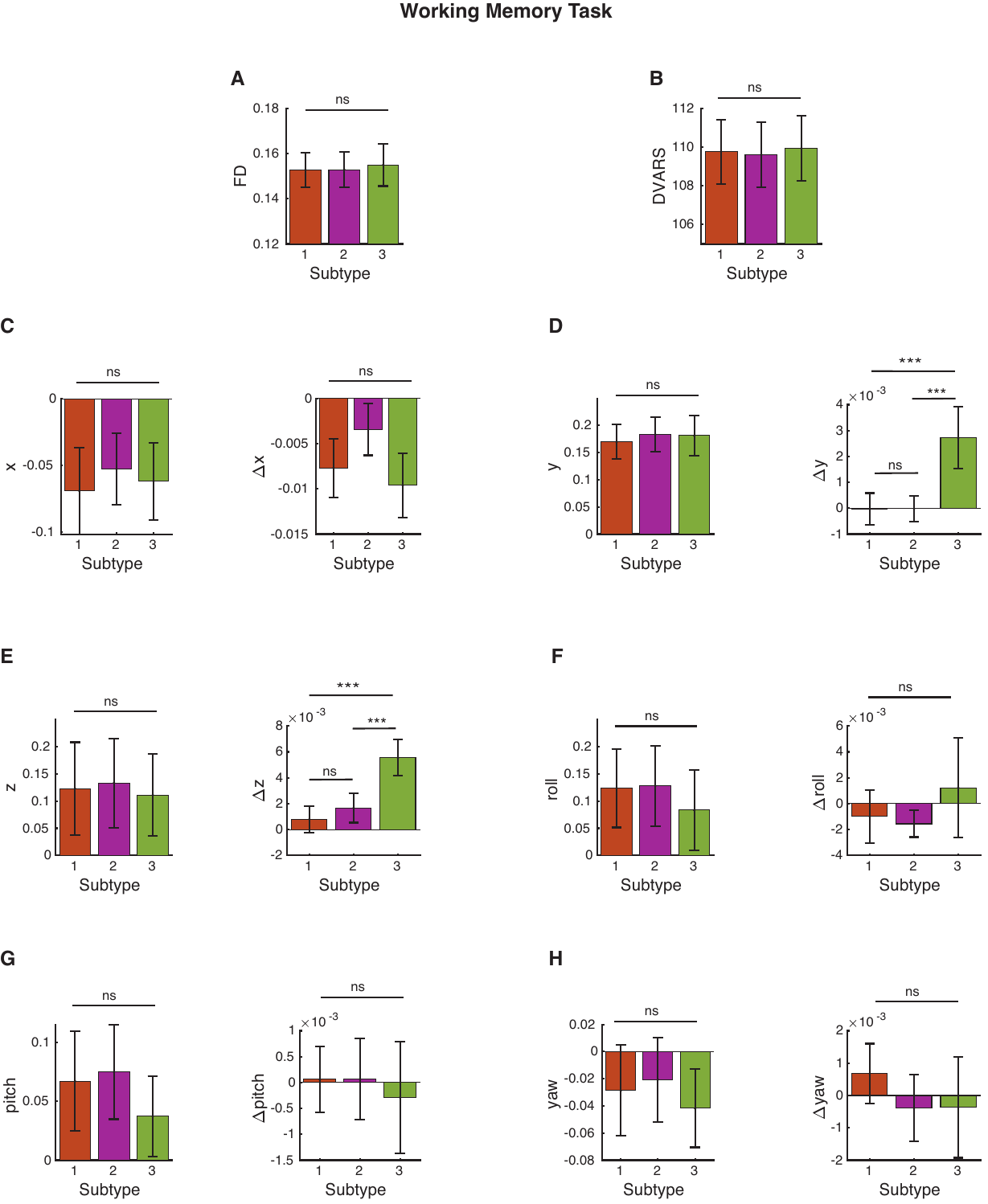

**Figure S15. Differences in motion parameters for the working memory task**. (A) Frame displacement (FD), (B) DVARS, (C) x-, (D) y-, (E) z-, (F) roll-, (G) pitch-, (H) yaw-direction. For each trial we estimated 14 different motion associated artifacts. Estimated motion values were averaged per subtype within a subject and statistical differences were determined using paired-samples t-test. For panels C-H, right panels show the 1^st^ derivatives. ***, P < 0.001; ns, not significant.
